# Deregulating m6A regulators leads to altered RNA biology in glioma cell lines

**DOI:** 10.1101/2024.10.28.620763

**Authors:** Syeda Maheen Batool, Hanna Lee, Ana K. Escobedo, Denalda Gashi, Kesli Faber, Prerna Khanna, Kase D. Haas, Tiffaney Hsia, Bob S. Carter, Leonora Balaj

**Affiliations:** Department of Neurosurgery, Massachusetts General Hospital, Harvard Medical School Boston, MA, USA

**Author notes:** Correspondence, L. Balaj, 185 Cambridge St. Boston, MA. Co-first authors (Syeda Maheen Batool, Hanna Lee, Ana K. Escobedo). Current address. University of Edinburgh, Medical School, Edinburgh, UK. University of Massachusetts, Chan Medical School, Worcester, MA, USA. University of Utah Health, Salt Lake City, UT, USA.

**Keywords:** N6-methyladenosine (m6A), RNA, epitranscriptomics, isoform, alternative splicing, signaling, glioma

## Abstract

N6-methyladenosine (m6A) is the most prevalent internal mRNA modification, enriched in the CNS yet poorly characterized in glioma. Using long-read RNA sequencing, we mapped m6A in an *in vitro* glioma model following knockdown (KD) of the reader IGF2BP2, writer METTL3, and eraser ALKBH5, with naive glioma cells and astrocytes as controls. Glioma cells exhibited a two-fold reduction in global m6A, suggesting progressive loss from healthy to malignant states. Integrated analysis revealed that m6A mediated control of gene expression is influenced by modification topology (CDS:3′UTR), transcript biotype, and length. Regulator KD, particularly ALKBH5 induced redistribution of m6A toward 3′UTR with consequent gene upregulation. We also identified m6A-mediated isoform switching, with a higher usage of retained intron and nonsense-mediated decay isoforms. Structural and splicing alterations at the isoform level were identified unique to each KD condition indicating m6A driven aberrant alternative splicing. At the functional level, KD specific remodeling of oncogenic signaling was also observed. ALKBH5 KD suppressed MYC targets and pro-apoptotic signaling while METTL3 KD enhanced mTOR and PI3K-AKT signaling. Collectively, these results demonstrate that m6A mediated regulation in glioma is highly context-dependent, defining distinct clinically relevant phenotypes. This has implications for future biomarker discovery and development of targeted therapeutics.

## Introduction

Over 163 RNA modifications have been identified, with N6-methyladenosine (m6A) the most prevalent modification detected in diverse RNA types (mRNA, tRNA, rRNA, circRNA, miRNA, lncRNA)^1,2^. m6A modification is dynamically regulated by three distinct groups of binding proteins, collectively termed m6A regulators: writers (methyltransferases), readers (m6A-binding proteins), and erasers (demethylases) which work together to maintain a dynamic balance in RNA modication^13^. The regulators influence multiple aspects of RNA metabolism, such as translation, alternative splicing, intracellular and extracellular transport, localization, and post-transcriptional stability^3,4^.

The role of m6A has been widely studied in physiological and pathological contexts, including malignancies of hematopoietic, reproductive systems and central nervous system^1,5^. Recent studies suggest that m6A is more enriched in CNS (central nervous system) relative to other organs^5^. Glioma is the most common primary CNS tumor, with glioblastoma being the most aggressive subtype^6^. Tumor recurrence and treatment resistance contribute to the poor prognosis of these heterogeneous lesions, with a median survival of approximately 15 months^6,7,8^. Emerging evidence implicates m6A in glioma progression^9^ through mechanisms such as enhancement of cell stemness^10^, tumor cell proliferation^11^, and modulation of the tumor microenvironment^12^. Conversely, other studies have demonstrated anticancer effects of m6A regulators, with lower m6A levels promoting tumorigenesis of glioma stem cells (GSCs)^13^. Together, these findings are suggestive of a potential dual, context-dependent role of m6A in glioma, underscoring the need for a more detailed exploration of its molecular mechanisms.

In this study, we sought to elucidate the precise molecular role of varying m6A levels in a glioma, using a Gli36 cell line model. We generated knockdown (KD) models for three key m6A regulators: IGF2BP2 KD (reader), METTL3 KD (writer), and ALKBH5 KD (eraser). Using Oxford Nanopore Technologies (ONT) long read direct RNA sequencing, we mapped the m6A-modified sites in RNA at single nucleotide resolution level. Each sample was sequenced in three independent runs to ensure reproducibility. Our findings provide new insights into the biological role of the epitranscriptomic landscape in glioma and may inform the development of targeted therapeutics.

## Results

### Quality assessment and transcriptomic analysis of m6A regulator knockdowns in glioma cells

We developed an *in vitro* glioma model with targeted knockdown (KD) of three key m6A regulators using small interfering RNAs (siRNAs): IGF2BP2 (reader), METTL3 (writer), and ALKBH5 (eraser) (Figure 1a, see Methods). As controls, we included untreated glioma cells (naive) and a non-diseased astrocyte cell line. Together, five experimental conditions of glioma cells were analyzed: (1) Naïve, (2) IGF2BP2 KD, (3) METTL3 KD, (4) ALKBH5 KD, and (5) normal astrocytes (see Methods). Direct RNA sequencing was performed for each condition in triplicate to ensure reproducibility (see Methods, Figure 1a).

**Figure 1.**
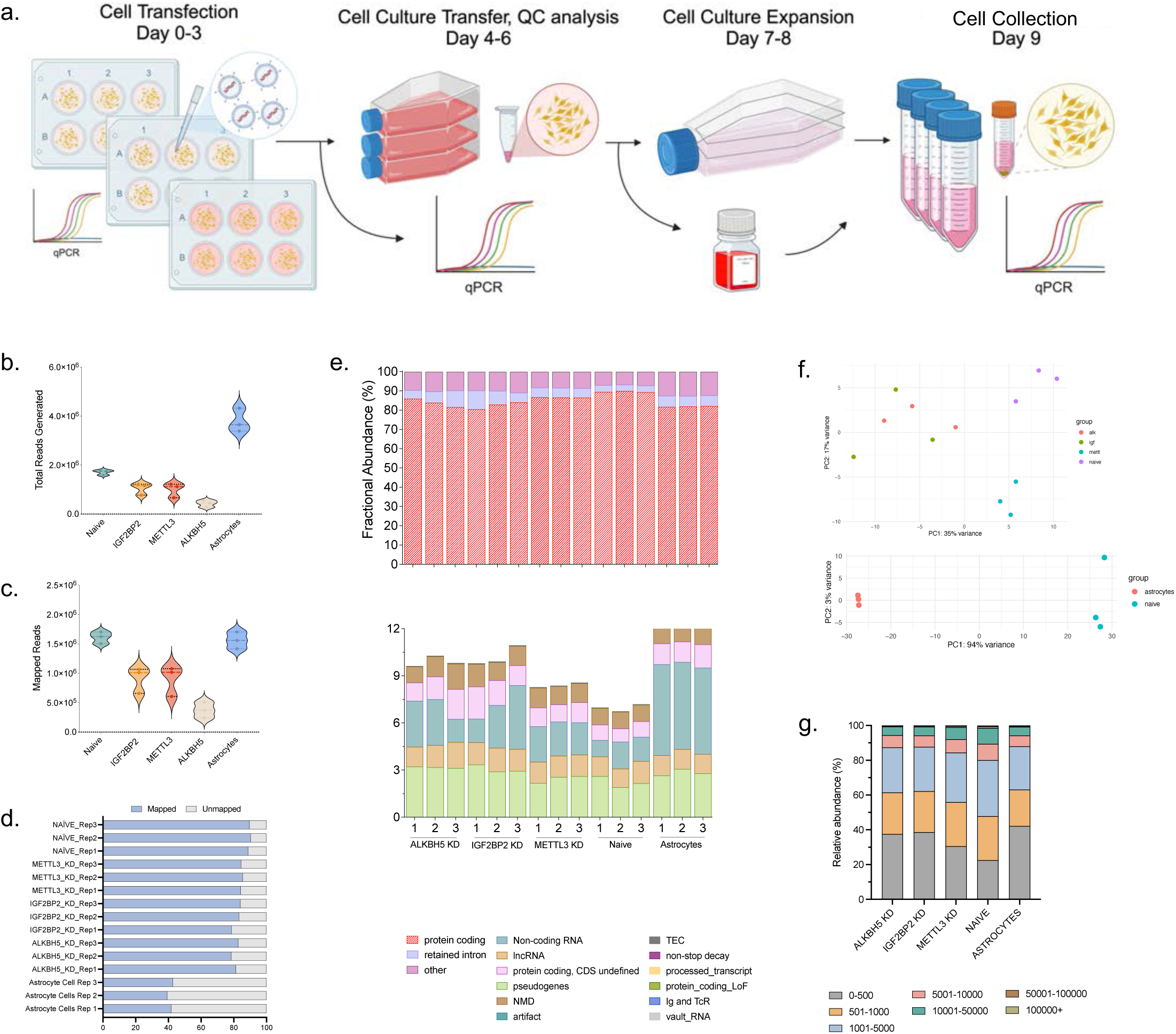
Transcriptome wide profiling m6A in glioma cell line. (a) Schematic depicting cell transfection with silencer RNA (siRNA) to achieve knockdown (KD) of key m^6^A regulators: IGF2BP2 (reader), METTL3 (writer), ALKBH5 (eraser). RNA isolated from control (naïve, astrocytes) and KD cells on day 9 sequenced using Nanopore direct RNA sequencing platform. Three technical replicates were sequenced per condition. (b) Total and (c) Mapped reads across the three replicates of cell RNA sequenced from glioma and KD conditions. (d) Assessment of percentage mapped vs. unmapped reads (e) Distribution analysis of the detected RNA biotypes from control and KD conditions. (f) Assessment of variance in the cell RNA detected transcripts using principal component analysis (PCA). (g) Distribution analysis of the detected RNA biotypes from control and KD conditions.

A total of 12.6 million reads were generated across all conditions (Figure 1b). While there was some variability in raw read counts, astrocytes yielded the highest number of reads (3,788,324), followed by the naive and knockdown (KD) conditions. A similar trend was observed in mapped reads (Figure 1c). Despite the variability in raw read counts, mapping efficiency remained consistent across conditions, with 80.9–84.8% of reads mapping in the KD conditions and 89% in the naive, untreated cells (Figure 1d). Interestingly, while astrocytes exhibited the highest number of raw and mapped reads, their mapping percentage was comparatively lower (42.9%) than that of naive and KD conditions (Figure 1d).

Protein-coding RNA was the predominant biotype across all conditions (Figure 1e). Naive glioma cells and METTL3 KD samples exhibited the highest proportions of protein-coding transcripts (Naive: 89.6%, METTL3 KD: 86.6%). In contrast, ALKBH5 KD, IGF2BP2 KD, and astrocytes exhibited modestly lower proportions (ALKBH5 KD: 83.8%, IGF2BP2 KD: 82.6%, Astrocytes: 82.0%). Retained intron RNA transcripts were more enriched in ALKBH5 KD (6.2%) and IGF2BP2 KD (7.3%) conditions, followed by astrocytes (5.6%) and METTL3 KD (5.0%), with naive glioma cells showing the lowest prevalence (3.4%) (Figure 1e).

Principal Component Analysis (PCA) was performed to assess global transcriptomic differences across conditions (Figure 1f). In the KD comparison (Figure 1f, top panel), PC1 and PC2 accounted for 35% and 17% of the variance, respectively. Naive glioma cells and METTL3 KD samples clustered closely, indicating minimal transcriptomic divergence. In contrast, IGF2BP2 KD and ALKBH5 KD samples formed discrete clusters, suggesting broader transcriptomic shifts upon KD of these regulators (Figure 1f, top panel). In the astrocyte-glioma comparison (Figure 1f, bottom panel), PC1 alone explained 94% of the variance, clearly segregating the astrocytes from glioma derived cells, reflecting their divergent transcriptomic landscapes (Figure 1f, bottom panel).

Analysis of the read length distribution demonstrated additional condition-specific patterns (Figure 1g). Shorter RNA transcripts (≤ 500 bp) were more frequently enriched in astrocytes (42.2%), followed by ALKBH5 KD (37.6%) and IGF2BP2 KD (38.7%) (Figure 1g). In contrast, longer (501-5,000 bp) RNA transcripts were more abundant in naive (57.4%) and METTL3 KD (53.8%) conditions, compared to astrocytes (45.8%) and the other KD conditions (ALKBH5 KD: 49.9%, IGF2BP2 KD: 49.0%) (Figure 1g).

Together, these preliminary results and quality control assessment highlight significant transcriptomic divergence following the KD of specific m6A regulators. Notably, IGF2BP2 and ALKBH5 KD cells exhibited distinct shifts, while METTL3 KD samples retained transcriptomic similarity to the naive condition. We also observed differences in RNA biotype composition and read length distribution, with ALKBH5 and IGF2BP2 KD cells showing higher enrichment of retained intron and shorter (≤ 500 bp) RNA transcripts. Conversely, METTL3 KD and naive glioma cells demonstrated a higher proportion of protein-coding RNA and longer (501-5,000 bp) transcripts.

### Transcriptome wide mapping of m6A in astrocytes, naive glioma cells and m6A regulator knockdowns

Having established global transcriptomic differences, we next examined the distribution and abundance of m6A modifications to determine how the regulator perturbations reshape the epitranscriptomic landscape. M6A RNA modifications were predicted using m6Anet (v2.1.0), a machine learning-based tool (see Methods). We quantified the proportion of modified sites, transcripts, and genes across all experimental conditions. Non-diseased astrocytes demonstrated the highest prevalence of m6A modifications, with 4.7% of sites, 33.8% of transcripts, and 38.3% of genes harboring at least one m6A mark (Figure 2a). In comparison, naïve glioma cells exhibited a significantly lower burden of m6A modifications, with only 2.0% of sites (2.3 fold lower), 23.4% of transcripts (1.4 fold lower), and 30.2% of genes (1.3 fold lower) showing evidence of m6A methylation (Figure 2a). All KD conditions displayed further reductions in m6A detection relative to both astrocytes and naive glioma cells. Among the regulators, ALKBH5 KD had the lowest m6A prevalence (1.2% of sites, 9.8% of transcripts, and 13.6% of genes), followed by METTL3 KD (15.1% of transcripts, 20.2% of genes) and IGF2BP2 KD (15.9% of transcripts, 21.4% of genes) (Figure 2a).

**Figure 2:**
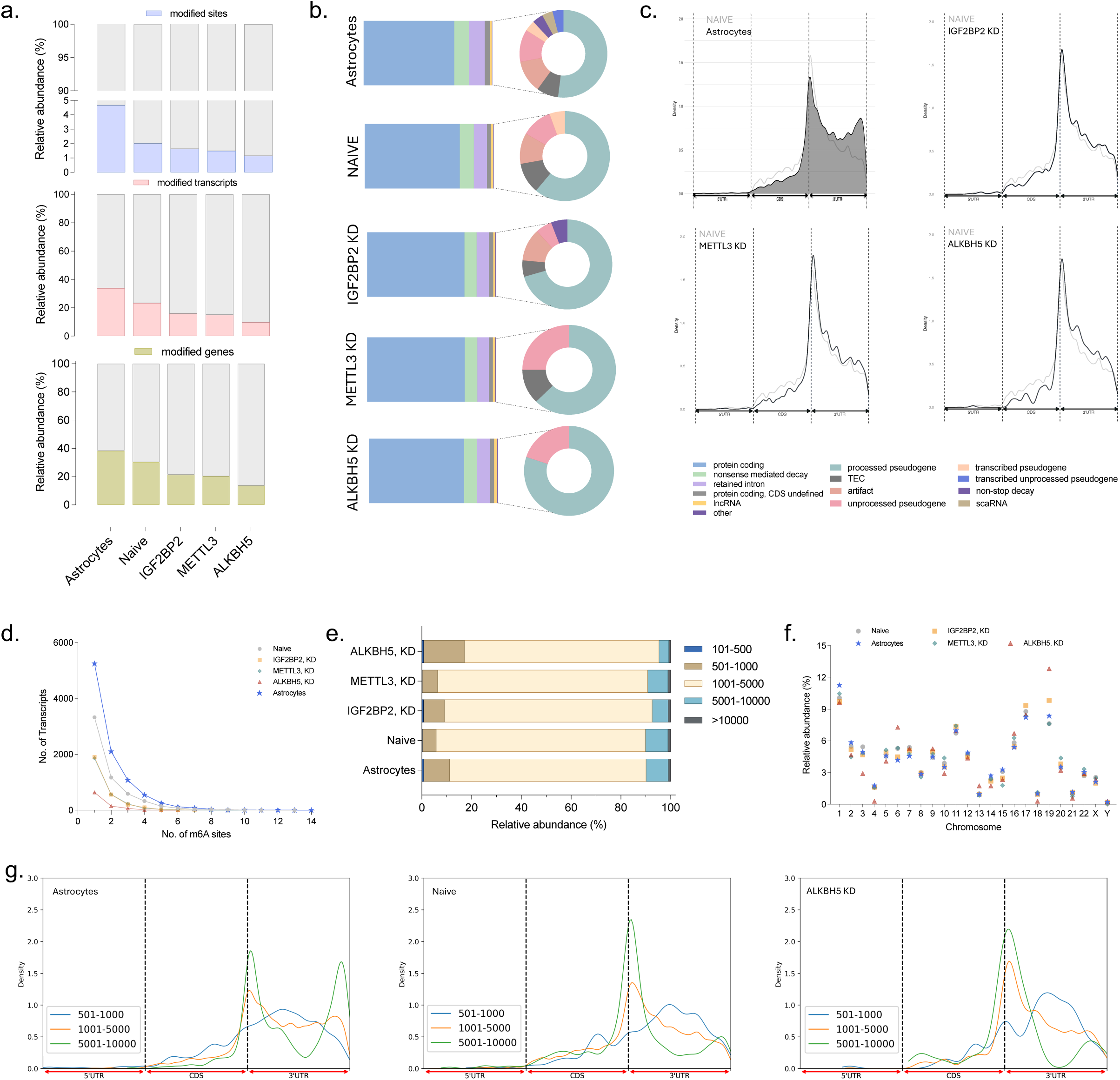
Quantification of m6A-modified RNA biotypes and transcript distribution. (a) Bar chart summarizing the relative abundance of m6A modified sites, transcripts, and genes for ALKBH5 KD, IGF2BP2 KD, METTL3 KD, naive, and astrocytes condition. (b) Pie chart depicting the m6a modified transcript biotype distribution in each condition (n=5). (c) KDE distribution of localization of m6A modified sites in the 5’UTR, CDS, and 3’UTR transcript regions in each KD condition (n=3) and Astrocytes (dark gray) in comparison to Naive (light gray). (d) Distribution of transcripts by number of m6A sites across experimental conditions, The y-axis indicates the number of transcripts, and the x-axis shows the number of m6A sites per transcript. (e) Stacked bar chart demonstrating the relative abundance of m6A modified RNA fragment length in each condition. (f) Scatter plot showcasing the chromosomal distribution of m6A modified genes. (g) Density plots of m6A distribution along the transcript region, stratified by RNA length.

No significant differences were observed among astrocytes, naive glioma cells and all KD conditions in the distribution of m6A-modified RNA biotypes. Protein-coding transcripts represented the majority of m6A-marked RNA (70.5%–75.5%), followed by transcripts undergoing nonsense-mediated decay (NMD, 9.4%–11.6%) and retained intron containing transcripts (9.0%–12.2%) (Figure 2b).

Transcriptomic mapping revealed condition-specific differences in m6A regional localization and enrichment patterns (Figure 2c). Astrocyte derived RNA was enriched for m6A in the 3′ untranslated region (3′UTR), whereas naive glioma cells exhibited greater m6A abundance in the coding sequence (CDS) region (Figure 2c). Notably, this inverse pattern was similarly observed across all KD conditions. Disruption of the m6A regulation via KD produced a consistent redistribution of m6A sites, characterized by a relative depletion in the CDS and increased localization to the 3′UTR. To control for bias, each KD condition was directly compared to naive, untreated glioma cells (Figure 2c). Among the KD groups, this shift was most pronounced in the ALKBH5 KD condition, where the loss of CDS-localized m6A and corresponding gain in 3′UTR methylation was particularly marked (Figure 2c).

We next examined the prevalence of single-versus multi-methylated transcripts (Figure 2d). A high proportion of transcripts contained only a single m6A site across all conditions: astrocytes (n = 5,251, 55%), naive (n = 3,324, 58%), IGF2BP2 KD (n = 1,899, 68%), METTL3 KD (n = 1,861, 67%), and ALKBH5 KD (n = 636, 74%) (Figure 2d). Astrocytes, with their higher overall m6A abundance, also had the greatest proportion of multi-methylated transcripts, with up to 14 sites per transcript (Figure 2d). In contrast, naive and all KD conditions exhibited a range in 6-10 m6A sites per transcript (Figure 2d).

Because m6A localization is linked to structural features of RNA, we assessed modification patterns across varying transcript length categories (Figure 2e). Regardless of cell type or m6A regulator KD, the highest prevalence of m6A modification was observed in transcripts ranging from >1,000 to ≤5,000 base pairs (78.2%–84.3%, Figure 2e). Short transcripts (≤500 bp), despite being highly abundant (Figure 1g), were rarely m6A methylated. Notably, astrocytes and ALKBH5 KD cells exhibited a higher proportion of m6A-modified transcripts in the 500–1,000 bp range (astrocytes: 10.4%; ALKBH5 KD: 16.4%) compared to naive (5.6%) and other KD conditions (IGF2BP2 KD: 8.4%; METTL3 KD: 6.2%). Transcripts 5,001–10,000 bp were infrequently methylated (<10% across all groups), with the lowest prevalence in ALKBH5 KD (3.9%) (Figure 2e). CDS and 3′UTR length comparisons revealed a pronounced shift in ALKBH5 KD cells, with shorter and less dense CDS regions, while 3′UTR lengths remained largely unchanged (Supplementary Figure 3a).

Chromosomal distribution analysis of m6A modified genes showed broadly similar enrichment patterns across astrocytes and glioma derived cell lines. The highest abundance of m6A modified genes was observed on chromosomes 1 (9.6%–11.3%), 17 (8.2%–9.3%), and 19 (7.6%–12.3%) (Figure 2f). Interestingly, chromosome 19 showed some divergence, with the highest proportion of m6A modified genes detected in the ALKBH5 KD condition (12.8%, Figure 2f).

Given the variation noted in m6A RNA length distribution, we also analyzed the transcriptomic distribution of m6A sites in RNA stratified by transcript length (Figure 2g). In astrocytes, m6A enrichment demonstrated prominent peaks at the CDS-3′UTR junction and within the tail end of 3′UTR, particularly in longer transcripts (>5,000bp). Naive glioma cells had similar enrichment near the CDS-3’UTR but reduced density within the 3′UTR (Figure 2g). ALKBH5 KD resulted in a broad loss of CDS enrichment and a flattened 3′UTR peak, indicating impaired regional specificity (Figure 2g). METTL3 KD, caused a dramatic reduction in global m6A density, particularly in long transcripts (Supplementary Figure 3c). In contrast, IGF2BP2 KD preserved some 3′UTR enrichment but showed an overall decrease in methylation (Supplementary Figure 3c).

In summary, these analyses revealed consistent differences in m6A RNA methylation across astrocytes, naïve glioma cells, and regulator knockdown conditions. Astrocytes exhibited the highest overall m6A prevalence, greater multi-site methylation, and predominant 3′UTR enrichment. Naive glioma cells and KD conditions showed progressively lower m6A levels, with KDs driving redistribution from CDS to 3′UTR. Transcript length and chromosomal localization also influenced modification likelihood, with the highest frequency observed in transcripts 1,000–5,000 bp in length and on chromosomes 1, 17, and 19. These features define the transcriptome-wide m6A landscape under healthy and glioma states, and reveal regulator-specific alterations in methylation patterns.

### Impact of m6A methylation on gene expression across pairwise comparisons

An important facet of m6A methylation as a regulatory system lies in its influence on post-transcriptional gene expression through context-dependent effects on RNA metabolism, including pre-mRNA processing, nuclear export, decay, and translation. To investigate this, we analyzed condition-specific patterns of m6A transcriptomic localization and their impact on gene expression across multiple pairwise comparisons.

We focused our analysis on differential methylation of 719 sites (578 transcripts, 244 genes) commonly detected (Supplementary Figure 3b) in naive and KD glioma cells (Figure 3a) (Supplementary Figure 4). m6A methylation was quantified per transcript by summing the m6A modification ratios of all detected sites per transcript, then normalizing by the transcript length to calculate a weighted modification ratio (see Methods). Hypermethylated (Log2FC > 0) or hypomethylated (Log2FC < 0) transcripts were subsequently correlated with gene expression levels (Figure 3b) (Supplementary Figure 4a). Six pairwise comparisons were determined: ALKBH5 KD vs Naive, IGF2BP2 KD vs Naive, METTL3 KD vs Naive, ALKBH5 KD vs IGF2BP2 KD, ALKBH5 KD vs METTL3 KD, IGF2BP2 KD vs METTL3 KD (Figure 3b).

**Figure 3:**
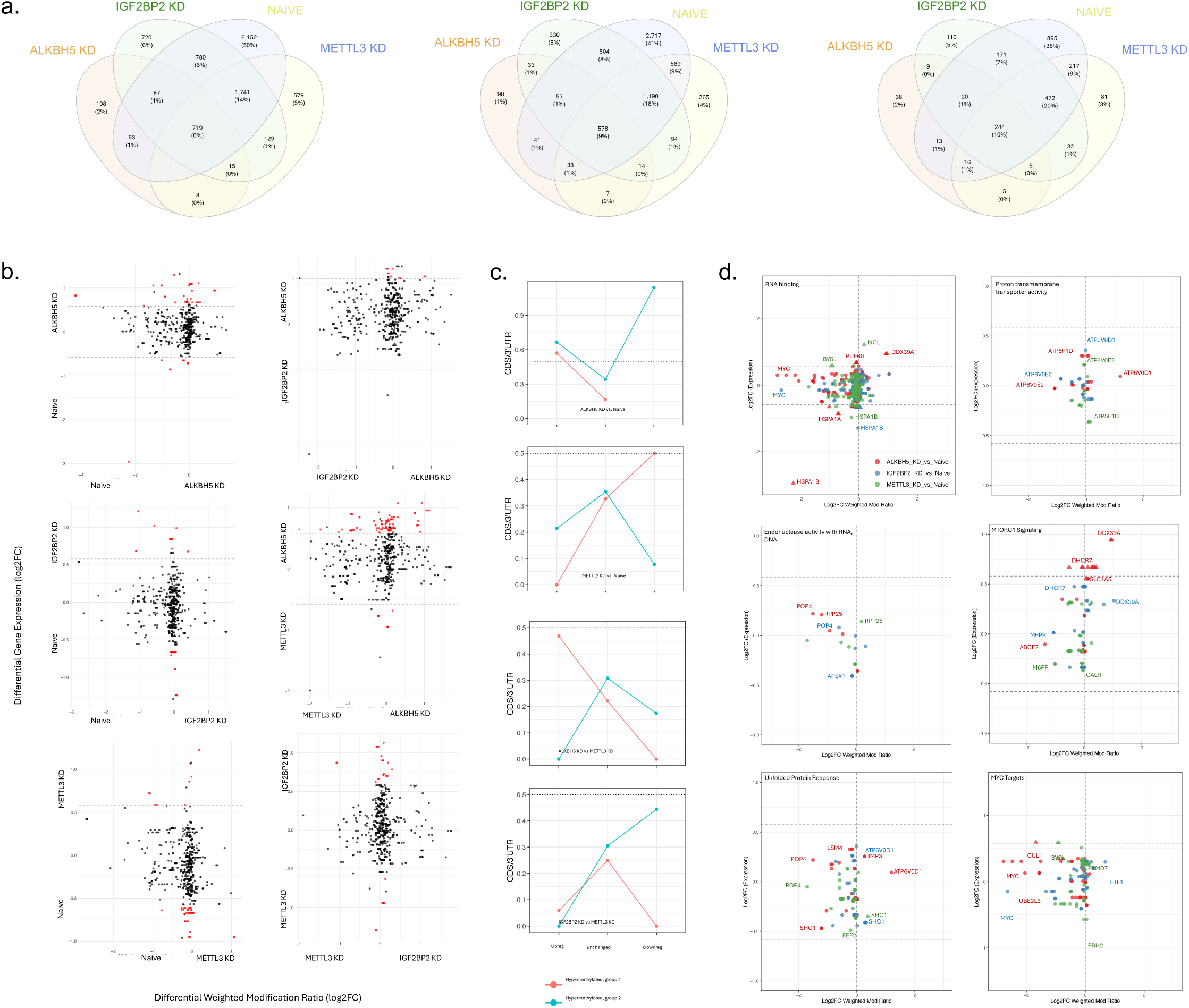
Relationship between m6A methylation, gene expression, transcript regions, and pathways across ALKBH5 KD, IGF2BP2 KD, METTL3 KD, and Naive conditions. (a) Venn diagram of the common and unique m6A modified sites, transcripts, and genes across the four conditions, respectively. N = 578 transcripts are used for downstream analysis. (b) Scatter plots illustrating the m6A methylation vs. differential gene expression of commonly m6A modified genes. The x-axis represents the log2 fold change (FC) in m6A methylation levels, while the y-axis denotes the log2 FC of differentially expressed genes (DGEs). DGEs with Log2 FC > 0.58 or < −0.58 and p-value < .05 are highlighted in red, while non-significant ones are shown in gray. (d) Line graph showcasing the ratio of abundance of CDS over 3’UTR in commonly modified m6A genes highlighting the relationship with gene expression and methylation in each knockdown comparison. (d) Scatter plots of selected pathways containing commonly m6A modified genes and displaying differential methylation and gene expression levels between in each knockdown condition compared to naive.

In comparison of naive to KD conditions, most commonly modified transcripts hypermethylated in the naive state (Figure 3b, left) (Supplementary Figure 4a). However, differential transcript-level methylation did not consistently translate into significant gene expression changes (Figure 3b, left). The extent of hypermethylation in naive cells varied by regulator KD and reflected the expression level of the targeted m6A regulator: (i) ALKBH5 KD (n = 23) vs Naive (n = 554), (ii) IGF2BP2 KD (n=15) vs Naive (n=562), (iii) METTL3 KD (n = 8) vs Naive (n = 569) (Figure 3b, left). ALKBH5 KD showed the most pronounced methylation loss (Log2FC ≥ –4), followed by IGF2BP2 KD (Log2FC ≥ –3), and METTL3 KD (Log2FC ≥ –2) (Figure 3b). Differential gene expression was only observed in ALKBH5 KD (upregulation in ALKBH5 KD), and METTL3 KD (upregulation in naive) but not IGF2BP2 KD (Figure 3b, left). Comparative analysis of the KD conditions revealed distinct clusters of hypermethylated transcripts unique to each group (Figure 3b, right). The largest methylation differences occurred in the Naive vs. ALKBH5 KD comparison. While ALKBH5 KD and IGF2BP2 KD showed no significant expression differences, ALKBH5 KD displayed more upregulated genes than both IGF2BP2 KD and METTL3 KD (Figure 3b, right).

To further assess the relationship between methylation and gene expression, we mapped m6A sites across transcript regions, stratifying transcripts by associated gene expression change: upregulated, unchanged, downregulated (Figure 3c) (Supplementary Figure 4b-d). In ALKBH5 KD vs. naive cells, higher m6A density in the CDS was associated with gene upregulation in naive cells, while lower CDS methylation corresponded with downregulation. Conversely, gene upregulation in ALKBH5 KD was more frequently linked to increased 3′UTR methylation. A similar trend was observed in METTL3 KD vs. naive cells: transcripts with exclusive 3′UTR methylation were upregulated in METTL3 KD, while m6A redistribution toward the CDS was linked to downregulation. Across all comparisons, naive glioma cells consistently exhibited a higher CDS/3′UTR methylation ratio among upregulated genes (Figure 3c). Within KD conditions, ALKBH5 KD showed gene upregulation associated with higher CDS methylation, whereas METTL3 KD demonstrated upregulation predominantly in transcripts with exclusive 3′UTR methylation (Figure 3c).

Given the context-dependent nature of m6A mediated regulation, we performed gene set enrichment analysis (GSEA) on commonly modified genes to identify enriched functional pathways (Figure 3d) (Supplementary Figure 5a). Integration of transcript-level weighted methylation ratios (x-axis) with gene expression changes (y-axis) across KD vs naïve comparisons revealed regulator-specific signatures (Figure 3d) (Supplementary Figure 4e).

In RNA binding pathways, ALKBH5 KD showed prominent hypomethylation, with upregulation of PUF60 and DDX39A, and downregulation of HSPA1A/B when compared to naive cells (Figure 3d). The MYC gene was consistently hypomethylated in both ALKBH5 and IGF2BP2 KD but was not differentially expressed (Figure 3d). Genes involved in RNA/DNA endonuclease activity, such as POP4 and RPP25, were broadly hypomethylated in ALKBH5 and IGF2BP2 KD, while RPP25 was hypermethylated in METTL3 KD (Figure 3d).

In proton transmembrane transport, ATP6V0E2 was hypomethylated in ALKBH5 and IGF2BP2 KD, while ATP6V0D1 was hypermethylated in ALKBH5 KD (Figure 3d). While most unfolded protein response genes were hypomethylated in KD cells, ALKBH5 KD uniquely exhibited both hypermethylation (IMP3, ATP6V0D1) and hypomethylation (POP4, SHC1) (Figure 3d). Similar methylation profiles were observed for ATP6V0D1 and SHC1 in IGF2BP2 KD cells (Figure 3d). In mTORC1 signaling, shared hypomethylation was observed for M6PR (METTL3 and IGF2BP2 KD) and DHCRP (ALKBH5 and IGF2BP2 KD)(Figure 3d). Notably, several genes were upregulated in ALKBH5 KD despite lacking differential methylation (Figure 3d).

Together, these analyses demonstrate that m6A methylation changes are highly regulator specific, with ALKBH5 KD producing the most pronounced loss of methylation and largest shift in regional distribution. The relationship between m6A deposition and gene expression was context-dependent, with CDS methylation generally linked to upregulation in naïve cells and 3′UTR methylation more frequently associated with upregulation in KD conditions.

### Unique m6A methylation landscapes in astrocytes, glioma, and regulator knockdowns

We next analyzed m6A sites, transcripts, and genes unique to astrocytes, naive glioma cells and each KD condition. Astrocytes harbored the highest number of unique m6A sites (11,967 sites, 5,327 transcripts, 1,458 genes), followed by naive glioma cells (6,148 sites, 3,653 transcripts, 1,520 genes), IGF2BP2 KD (720 sites, 682 transcripts, 355 genes), METTL3 KD (579 sites, 536 transcripts, 281 genes), and ALKBH5 KD (200 sites, 194 transcripts, 104 genes) (Figure 4a).

**Figure 4:**
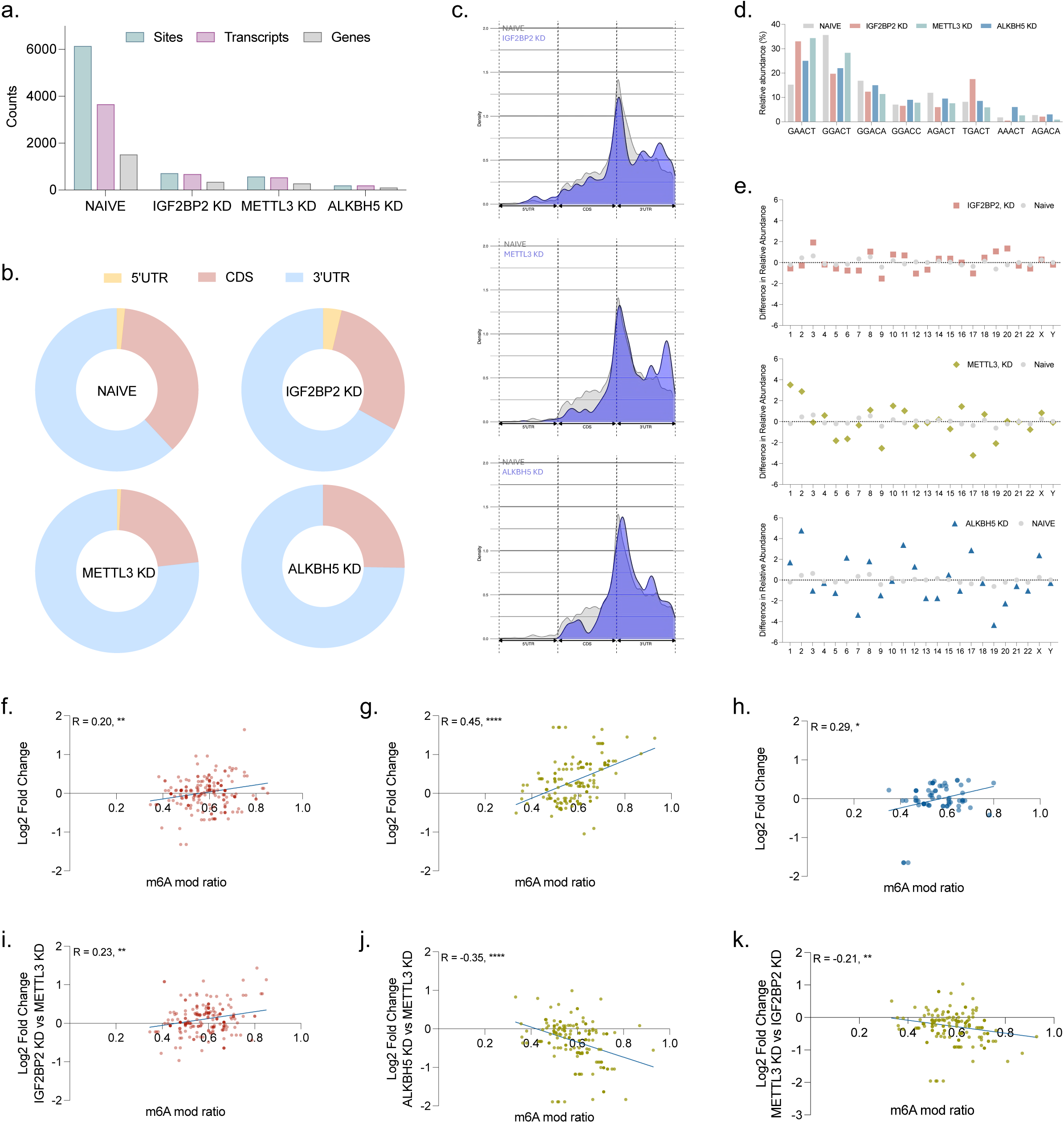
Quantification of unique m6A modified sites, transcripts, and genes. (a) Grouped bar chart displaying the number of unique m6A modified sites, transcripts, and genes in each KD cell and the naive condition. (b) Pie charts representing the proportion of 5’UTR, CDS, and 3’UTR in uniquely modified m6A transcripts for each KD condition (n=3) and the naive condition. (c) Line graphs showing the transcript region density in ALKBH5 knockdown, IGF2BP2 knockdown, and METTL3 knockdown (blue) in comparison to naive (gray). (d) k-mer distribution in each knockdown comparison and naive.(e) Scatter plot depicting the chromosomal distribution in each m6A regulator knockdown when compared to naive. (f-k) Scatter plots of the m6A modified ratio (x-axis) against the log2 fold change to showcase the correlation between methylation and gene expression. Yellow represents METTL3 knockdown, red represents IGF2BP2 knockdown, and blue represents ALKBH5 knockdown.

Biotype distributions of uniquely modified m6A transcripts were broadly similar across conditions (Supplementary Figure 5b). Protein coding m6A RNA transcripts were more prevalent in ALKBH5 KD (77.3%) and IGF2BP2 KD (75.1%), compared to naive (73.3%) and METTL3 KD (73.1%). Among non-coding RNAs, METTL3 KD exhibited the highest proportion of m6A modified lncRNAs (3.4%) and NMD-targeted transcripts (13.4%) (Supplementary Figure 5b).

Transcriptomic regional mapping revealed condition specific m6A localization patterns (Figure 4b-c). Within KD cells, 5’UTR modifications were detected only in IGF2BP2 KD (3.7%), a 2.3-fold higher enrichment compared to naive glioma cells (1.6%) (Figure 4b-c). Consistent with the common site analysis, naive cells exhibited the highest CDS (36.4%) and lowest 3’UTR (62.0%) methylation, yielding the highest CDS:3’UTR ratio (0.59). Similar to the earlier findings on m6A mediated gene expression regulation, all KD conditions showed reduced CDS methylation and increased 3′UTR methylation (Figure 4b-c). All KD conditions showed reduced CDS methylation and increased 3′UTR methylation: ALKBH5 KD (CDS: 25.3%, 3′UTR: 74.7%), METTL3 KD (22.4%, 76.8%), and IGF2BP2 KD (29.4%, 67.0%) (Figure 4b-c). Overall, CDS:3′UTR ratios were lower in all KD conditions relative to naive, with METTL3 KD showing the lowest ratio (0.29), followed by ALKBH5 KD (0.33) and IGF2BP2 KD (0.44) (Figure 4b–c).

K-mer analysis of unique m6A sites revealed uneven distribution across conditions (Figure 4d). Consistent with previous findings, GGACT was the most enriched motif in naive cells (35.7%). However, KD of the distinctive m6A regulators led to downregulation of GGACT and a 2 fold higher prevalence of GAACT (METTL3 KD: 34.4%, IGF2BP2 KD: 33.1%, ALKBH5 KD: 25.0%) compared to naive cells (Figure 4d). GGACT depletion was most pronounced in IGF2BP2 (19.7%) and ALKBH5 KD (22.0%) cells, with METTL3 KD retaining a relatively higher level (28.3%) (Figure 4d). Notably, TGACT was most enriched in IGF2BP2 KD (17.5%), while AGACT was more prevalent in naive cells (11.9%) compared to ALKBH5 (9.5%), METTL3 (7.6%), and IGF2BP2 KD (6.0%) (Figure 4d).

Earlier chromosomal distribution analysis of all detected m6A sites did not reveal condition specific genomic targeting. However, when the analysis was restricted to unique m6A sites, distinct chromosomal patterns emerged across KD conditions relative to the naive state (Figure 4e). To quantify this, we first computed the average chromosomal prevalence of m6A-modified transcripts in naïve cells, which served as the baseline for calculating the shift in each KD condition. The resulting “difference in relative abundance” was plotted to highlight deviations (Figure 4e). No significant shift was observed in IGF2BP2 KD compared to naive cells. In contrast, ALKBH5 KD, followed by METTL3 KD exhibited pronounced alterations in the chromosomal distribution of m6A modified genes (Figure 4e). Specifically, METTL3 KD was associated with an increased number of modified targets on chromosomes 1 and 2, and a reduction on chromosomes 9 and 17 (Figure 4e). ALKBH5 KD resulted in an upregulation of m6A-modified transcripts across multiple chromosomes, most notably chromosomes 2, 11, and 17 (Figure 4e).

To assess functional effects of these modifications, we performed a correlation analysis between the m6A weighted modification ratio (see Methods) and the corresponding gene expression levels (Figure 4f–k). In all KD conditions, uniquely modified transcripts showed a positive correlation with gene expression relative to naIve (Figure 4f–h). This correlation was strongest in METTL3 KD (R = 0.45), followed by ALKBH5 KD (R = 0.29) and IGF2BP2 KD (R = 0.20), with all correlations reaching statistical significance. Comparative analyses across KD conditions revealed that IGF2BP2 KD specific m6A modified transcripts were associated with higher gene expression in IGF2BP2 KD versus METTL3 KD (Figure 4i–j), whereas ALKBH5 KD specific transcripts were linked to downregulated gene expression in ALKBH5 KD relative to METTL3 KD (Figure 4k).

In summary, naive glioma cells demonstrated preferential CDS m6A localization with enrichment of GGACT. M6A regulator KD altered both the regional and chromosomal distribution of the m6A sites, with also a functional impact on gene expression. These findings underscore the divergent and regulator specific impacts of METTL3, IGF2BP2, and ALKBH5 in shaping m6A dynamics.

### Expression dynamics of m6A regulators, splicing factors, and decay machinery

We next examined how m6A regulator knockdown affected the expression of m6A-related genes and their isoforms across seven pairwise comparisons: ALKBH5 KD vs. naive, IGF2BP2 KD vs. naive, METTL3 KD vs. naive, ALKBH5 KD vs. IGF2BP2 KD, ALKBH5 KD vs. METTL3 KD, IGF2BP2 KD vs. METTL3 KD, and astrocytes vs. naïve (Figure 5a).

**Figure 5:**
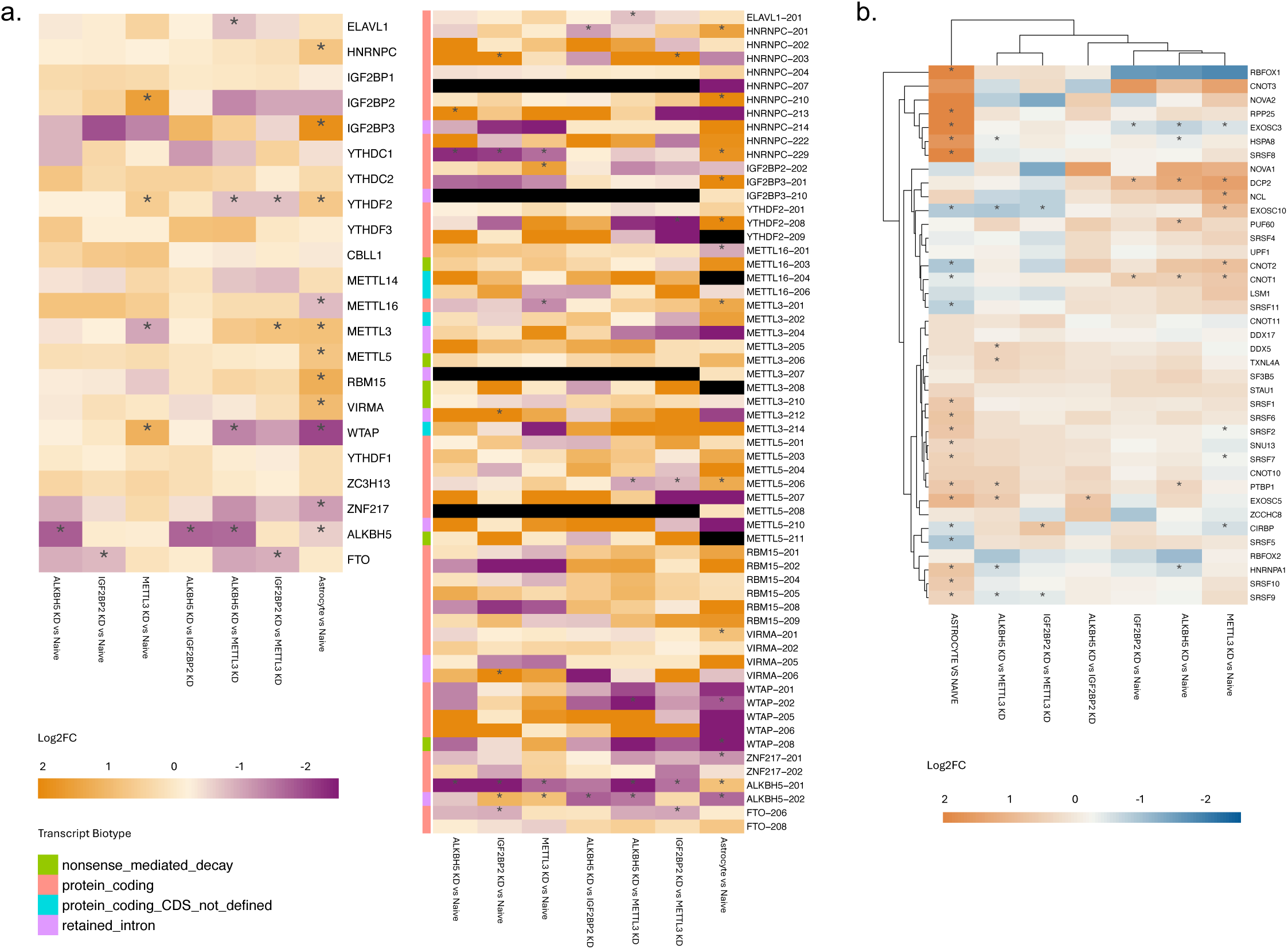
Distribution analysis of m6A regulators, RNA splicing and decay factors across naive and KD cellular states. (a) Heatmaps summarizing the differential gene (left) and transcript (right) expression for m6A regulators. Each row represents a specific m6A RNA regulator gene/transcript and the columns represent the log2 fold change of expression for the seven pairwise comparisons. For KD conditions vs Naïve, orange indicates gene/transcript upregulation in the KD condition and purple indicates upregulation in the Naïve condition. For Naïve vs Astrocytes, orange indicates upregulation in Naïve and purple indicates upregulation in Astrocytes. Significant differential expression is indicated with a star and transcript biotypes are provided in the row annotations. (b) Heatmap depicting RNA splicing and decay factors in relation with gene expression. For KD conditions vs Naïve, red indicates gene/transcript upregulation in the KD condition and blue indicates upregulation in the Naïve condition. For Naïve vs Astrocytes, red indicates upregulation in Naïve and blue indicates upregulation in Astrocytes. Genes with significant changes in expression are highlighted with a star.

As expected, ALKBH5 was significantly downregulated in all ALKBH5 KD comparisons (vs naive: −1.80; vs IGF2BP2 KD: −1.72; vs METTL3 KD: −1.66) and in astrocytes vs naive (−0.65). At the isoform level, ALKBH5-201 mirrored this trend in KD comparisons but was upregulated in astrocytes vs naive (0.80).

YTHDF2, a reader linked to RNA decay, was upregulated in METTL3 KD vs naive (0.55) but downregulated in ALKBH5 KD vs METTL3 KD (−0.77) and IGF2BP2 KD vs METTL3 KD (−0.79). It was also upregulated in astrocytes vs naïve (0.61). Its isoform YTHDF2-208 showed a similar pattern, with marked upregulation in astrocytes vs naive (3.53) and METTL3 KD vs naïve (1.80), and strong downregulation in ALKBH5 KD vs METTL3 KD (−2.26) and IGF2BP2 KD vs METTL3 KD (−3.49). Given that YTHDF2 can promote glioblastoma progression via PI3K/AKT activation, these data suggest that METTL3 KD may contribute to tumorigenic signaling (Figure 5a).

IGF2BP3 was significantly upregulated in astrocytes vs naive (1.84) and downregulated in all KD vs naive comparisons (ALKBH5 KD: −0.92; IGF2BP2 KD: −1.94; METTL3 KD: −1.39). Isoform IGF2BP3-201 followed the same trend, with reduced expression in all KD vs naive comparisons (−1.65 to −1.32) and elevated expression in astrocytes vs naive (2.13). In contrast, IGF2BP2 and IGF2BP2-202 were upregulated in all KD vs naive comparisons (IGF2BP2: 0.16 to 1.49; IGF2BP2-202: 0.11 to 1.48) (Figure 5a).

ZNF17 and its isoform ZNF17-201, both associated with reduced m6A levels, were significantly downregulated in astrocytes vs naive (−1.11) (Figure 5a). WTAP was downregulated in astrocytes vs naive (−2.04) but showed variable expression across KDs, including modest upregulation in METTL3 KD vs naïve (0.29) and downregulation in IGF2BP2 KD vs METTL3 KD (−0.78) and ALKBH5 KD vs METTL3 KD (−1.26). Isoforms WTAP-201, WTAP-202, and WTAP-208 followed a similar pattern, with significant downregulation in astrocytes vs naive (−1.84 to −2.80) and selective upregulation in METTL3 KD comparisons (Figure 5a).

Several splicing and decay factors exhibited condition-specific regulation (Figure 5b). In METTL3 KD vs naive, we observed downregulation of SRSF2 (−0.25), SRSF7 (−0.27), and EXOSC3 (−0.50), alongside upregulation of CNOT1 (0.61), CNOT2 (0.81), and EXOSC10 (0.81). Astrocytes vs naive displayed the opposite trend, with upregulation of SRSF2 (0.66), SRSF7 (0.46), and EXOSC3 (2.05), and downregulation of CNOT1 (−0.41), CNOT2 (−0.89), and EXOSC10 (−0.86) (Figure 5b).

All KD vs naive comparisons showed consistent upregulation of CNOT1 (0.49 to 0.61) and DCP2 (0.92 to 1.43) and downregulation of EXOSC3 (−0.50 to −0.72). The SRSF family was generally upregulated in astrocytes vs naïve, except SRSF5 (−0.88) and SRSF11 (−0.79). RBFOX1 was strongly downregulated in all KD vs naïve comparisons (−1.58 to −1.79) but showed a striking increase in astrocytes vs naïve (4.07) (Figure 5b).

Building on these observations, we next examined how these regulator-specific methylation and expression changes converge at the isoform level to reveal transcript structural alterations and functional consequences.

### Isoform switching and structural consequences of m6A regulator knockdown

Isoform switch analysis was performed using IsoformSwitchAnalyzeR^14^ (Supplementary Figure 6). The highest number of isoforms switching events was observed between astrocytes and naive glioma cells (2655), with comparable numbers of high-usage isoforms identified in each group (astrocytes: 1356; naive: 1299) (Figure 6a). Among the KD conditions relative to naive cells, ALKBH5 KD exhibited the highest number of isoform switches (1364), followed by IGF2BP2 KD (821) and METTL3 KD (459) (Figure 6a) (Supplementary Figure 6a-b).

**Figure 6:**
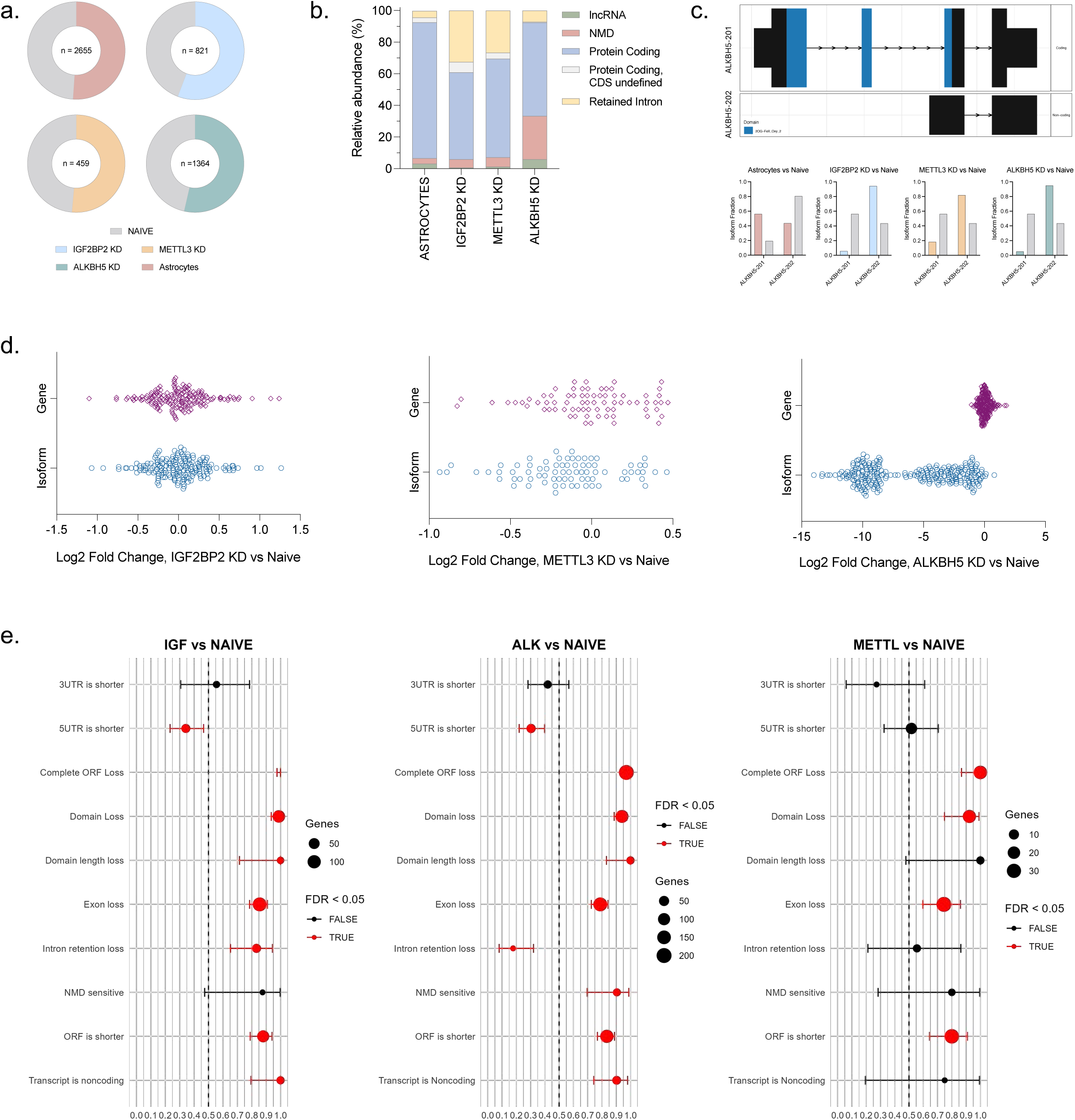
Quantification of Isoform Switching events and its correlation to gene and isoform expression. (a) Pie chart quantifying the isoforms switches occurring between each KD condition and astrocytes compared to naïve condition. (b) Stacked bar chart indicating the biotype distribution of isoforms that are used more in KDs and astrocytes compared to naive. (c) Isoform switching of ALKBH5-201 and ALKBH5-202 in astrocytes and KD when compared to naïve glioma cells. (d) Scatter plots of the log2 fold change at the isoform and the gene level of isoforms that are used more in the ALKBH5 knockdown, IGF2BP2 knockdown, and METTL3 knockdown. (e) Isoform switch consequence plots that highlight the effect an isoform causes for the knockdown conditions. The x-axis depicts the ratio of number of genes with the specific consequence over the total number of genes effected by that consequence with the y-axis specifying the consequence.

Notably, high-usage isoforms in astrocytes were predominantly protein coding (86.0%), whereas KD conditions showed a reduced proportion of protein coding isoforms, ranging from 55.0% to 62.3% (Figure 6b). Conversely, non-protein coding isoforms, including retained intron and nonsense mediated decay (NMD) targeted transcripts, were more frequently utilized in KD cells, consistent with a deregulated m6A machinery. ALKBH5 KD cells exhibited a markedly higher proportion of high-usage NMD transcripts (27.4%), approximately 5-fold greater than other KD conditions and 8-fold higher than astrocytes (Figure 6b). Additionally, IGF2BP2 and METTL3 KD cells displayed a significant enrichment of retained intron isoforms (IGF2BP2: 32.5%, METTL3: 26.7%), substantially exceeding the levels observed in ALKBH5 KD (7.1%) and astrocytes (4.2%) (Figure 6b) (Supplementary Figure 6b). An important example is the significant isoform switching observed for ALKBH5 when comparing KD cells and astrocytes to naive (Figure 6c). Interestingly, astrocytes showed higher usage of the protein-coding ALKBH5-201 isoform and lower usage of the non-coding ALKBH5-202 isoform compared to naive cells. In contrast, all KD conditions exhibited preferential usage of ALKBH5-202, whereas ALKBH5-201 was more abundant in naive cells (Figure 6c).

Differential usage of non-coding isoforms can impact gene expression. To explore this relationship, we plotted gene level and isoform level expression changes (Log2FC, KD vs Naive) for the high usage non-coding isoforms in each KD condition (Figure 6d). In IGF2BP2 KD cells, high usage of retained intron isoforms was associated with heterogeneous expression profiles, contributing to both upregulation and downregulation of genes and isoforms (Figure 6d). In contrast, METTL3 KD predominantly exhibited concurrent downregulation of gene and isoform expression associated with increased usage of retained intron isoforms (Figure 6d). The most striking pattern was observed in ALKBH5 KD cells, where high usage of nonsense mediated decay (NMD) isoforms resulted in minimal changes in gene-level expression but consistent and pronounced downregulation of the corresponding isoforms, with no instances of isoform upregulation (Figure 6d).

Structural consequence analysis identified features commonly altered across KD vs naive comparisons (Figure 6e). Several consequences were shared among all three KD conditions, including complete open reading frame (ORF) loss, domain loss, exon loss, and shorter ORFs. Nonsense mediated decay sensitive isoforms were uniquely detected in ALKBH5 KD cells (Figure 6e). Additional consequences shared between IGF2BP2 KD and ALKBH5 KD included shortened 5′UTRs, domain length loss, intron retention loss, and increased usage of noncoding transcripts (Figure 6e). Across all KD conditions, splicing events exhibited a general dysregulation when compared with naive cells: intron retention, ATSS (alternative transcription start site) gain, ATTS (alternative transcription termination site) gain, and increased usage of alternative 5′ and 3′ splice sites (A5, A3 gain) (Supplementary Figure 6c). We also performed gene ontology analysis to identify enriched pathways because of selected structural alterations (Supplementary Figure 7).

The results from isoform switching analysis reflect the underlying heterogeneity in isoform expression and usage, which in turn together mediate the associated gene expression. Knockdown of specific m6A regulators induced distinct shifts in isoform composition and structural features, underscoring the transcript level consequences of m6A mediated regulation.

### Functional consequences of m6a regulator knockdown on core oncogenic and survival-related pathways

To assess the functional consequences of m6A regulator knockdown, we examined both gene expression and m6A modification patterns across six core oncogenic and survival-related signaling cascades: apoptosis, MAPK, mTOR, oxidative Phosphorylation, PI3K-AKT, and MYC signaling (Figure 7a–f) (Supplementary Figure 8, 9). Hierarchical clustering of differentially expressed genes within each pathway revealed distinct expression patterns across KD conditions compared to naive glioma cells.

**Figure 7:**
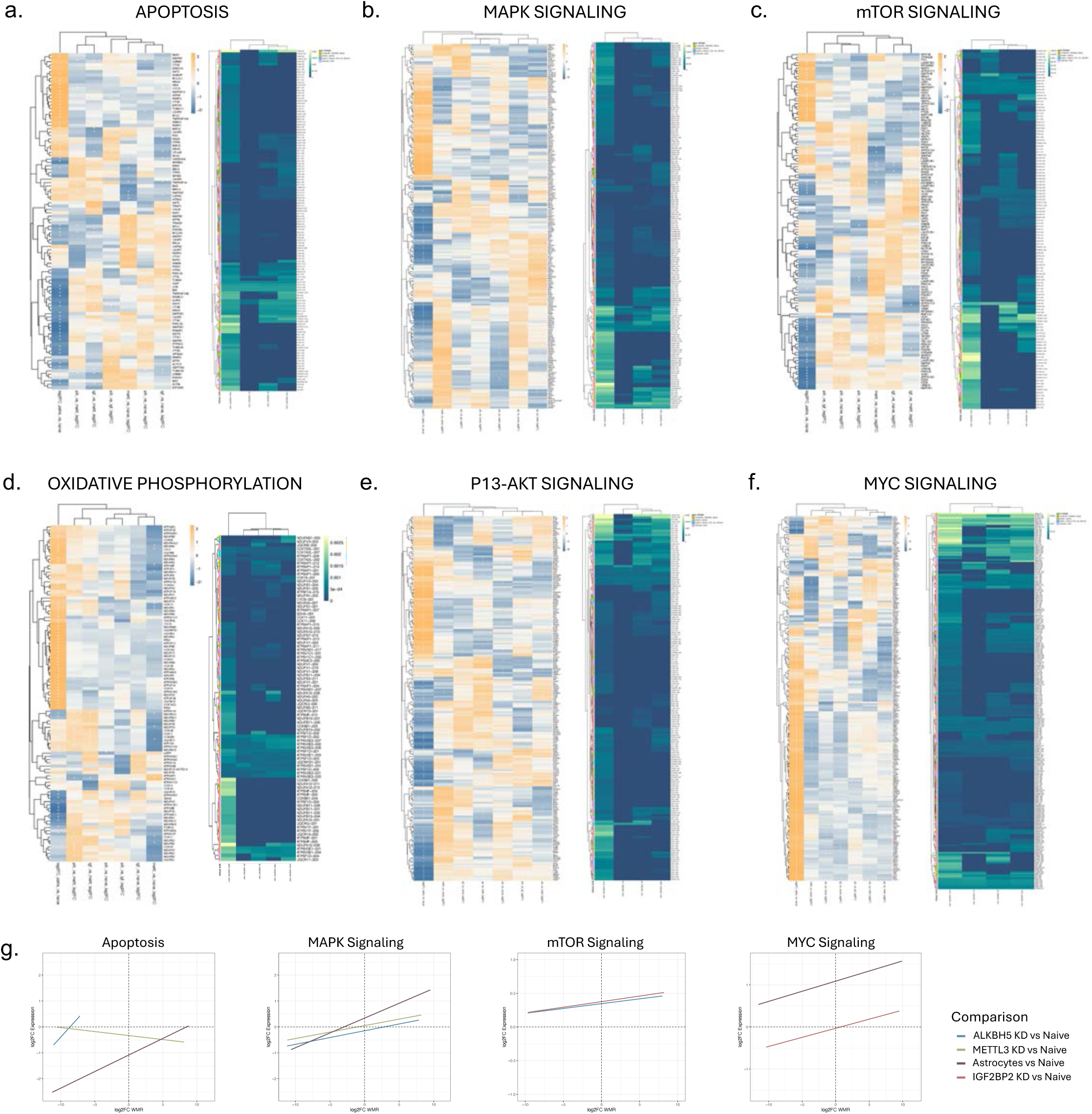
The relationship between selected genes in enriched pathways and the effect on gene expression and methylation. (a-f) Heatmaps (left) for selected significantly enriched gene ontology pathways where each row represents a gene involved in the specific pathway. The x-axis indicates the log2 fold change in gene expression between each group. Positive log2 fold change value indicate an upregulation in condition 1 and negative log2 fold change values indicate upregulation in condition 2. The heatmap (right) also displays the relation between selected genes and the m6A methylation levels in specified enriched pathways. (g) Correlation analysis between m6A methylation in transcripts and gene expression in Knockdown (KD) vs Naïve glioma cells in selected pathways.

Each pathway was analyzed by hierarchical clustering of differentially expressed genes and matched with heatmaps of corresponding m6A modification ratios (Supplementary Figure 8). Scatter plots of Log2 Fold Change in gene expression vs. transcript level methylation were generated to evaluate the correlation between these two layers in KD vs. naive comparison (Figure 7g).

ALKBH5 KD showed strong downregulation of pro-apoptotic genes (MAP2K2, BBC3/PUMA, CASP3) with parallel hypomethylation (Figure 7a, g), suggesting that reduced methylation may contribute to suppressed apoptotic priming. IGF2BP2 KD exhibited upregulation of extrinsic apoptotic effectors (CTSL, TNFRSF10B, DIABLO) accompanied by increased m6A levels, indicating a potential methylation-associated enhancement of apoptotic signaling. METTL3 KD had mixed patterns, with anti-apoptotic genes (BCL2A1, MCL1) being upregulated without major methylation shifts, while TP53 targets remained largely unchanged (Figure 7a, g).

In the MAPK signaling cascade (Figure 7b, g), IGF2BP2 KD showed coordinated upregulation and hypermethylation of MAPK1, DUSP6, and FOS, consistent with methylation-associated activation. ALKBH5 KD broadly suppressed MAPK members (MAP3K5, MAP2K4, JUN) with concurrent hypomethylation, suggesting dampened pathway signaling. METTL3 KD selectively upregulated ERK2/MAPK1 and ELK1, both showing modest methylation gains (Figure 7b, g).

In mTOR signaling (Figure 7c, g), ALKBH5 KD downregulated RHEB, RICTOR, and EIF4EBP1 alongside hypomethylation, consistent with suppressed translational control. METTL3 KD upregulated AKT1, MTOR, and RPTOR, each displaying hypermethylation, implying that m6A enrichment may support pathway activation. IGF2BP2 KD showed increased TSC1 and DEPTOR expression (negative regulators) with accompanying hypermethylation, suggesting upstream feedback inhibition (Figure 7c, g).

In oxidative phosphorylation (Figure 7d), IGF2BP2 KD exhibited pronounced downregulation and hypomethylation of electron transport chain genes (NDUFB6, COX6C, ATP5F1C), indicating mitochondrial impairment potentially linked to methylation loss. ALKBH5 KD displayed moderate suppression and hypomethylation of Complex I and IV subunits. METTL3 KD largely preserved OXPHOS expression and methylation, indicating minimal disruption to mitochondrial bioenergetics (Figure 7d).

P13-AKT signaling (Figure 7e) was seen to be impacted significantly upon m6A writer and eraser KD. METTL3 KD upregulated PIK3R1, AKT2, and GSK3B with corresponding hypermethylation, supporting a pro-survival methylation–expression coupling. ALKBH5 KD downregulated PDK1, AKT1, and FOXO3 with parallel hypomethylation, indicating attenuation of downstream signaling. IGF2BP2 KD displayed a mixed profile, with PTEN and GSK3A upregulated (hypermethylated) and AKT3 downregulated (hypomethylated), suggesting pathway rewiring toward growth inhibition (Figure 7e).

A robust suppression of MYC target genes (NPM1, CDK4, ODC1, CCND1) was observed in ALKBH5 KD with broad hypomethylation (Figure 7f, g), consistent with reduced proliferative signaling. METTL3 KD maintained expression and methylation of key MYC targets (EIF4E, CDK6), suggesting sustained pathway output. IGF2BP2 KD showed selective downregulation and hypomethylation of cell cycle regulators while retaining ribosomal biogenesis components with relatively unchanged methylation, indicating partial MYC pathway disruption (Figure 7f, g).

Across pathways, ALKBH5 KD was characterized by broad hypomethylation coupled to reduced expression in pro-growth and survival networks, METTL3 KD by hypermethylation associated with maintained or enhanced oncogenic signaling, and IGF2BP2 KD by mixed methylation-expression patterns that often reflected pathway inhibition. These findings highlight the distinct and context-dependent ways in which m6A regulators influence both the transcriptional and epitranscriptomic landscape of critical signaling axes in glioma.

## Discussion

In this study, we profiled m6A methylation in an *in vitro* glioma model and compared it to non-diseased astrocytes to test the hypothesis that perturbation of m6A regulatory machinery produces distinct and context-dependent transcriptomic, isoform, and pathway-level consequences. To systematically dissect these effects, we selectively silenced three key m6A regulators: IGF2BP2 (reader), METTL3 (writer), and ALKBH5 (eraser). Our integrative approach allowed us to resolve the downstream functional and biological consequences of m6A dysregulation in glioma.

Compared to astrocytes, naive glioma cells exhibited a 2-fold reduction in overall m6A prevalence, with high confidence modifications detected in only 2% of all detected sites, 23.4% of transcripts, and 30.2% of genes. Notably, these findings align with our previous observations in a glioma tumor tissue model^15^, where direct comparison of non-tumor cortex, IDH1-mutant glioma, and glioma tissue-derived RNA revealed significant differences in m6A site detection. The highest prevalence was observed in the entry cortex, followed by IDH1-mutant glioma, with the lowest number of sites detected in glioma tissue^15^. Collectively, these results from both align with the idea that normal, nondiseased CNS tissue harbors a high basal level of m6A modifications that may become progressively dysregulated in tumors, including glioma. Furthermore, the extent of methylation loss or dysregulation in gliomas appears to vary with the underlying molecular alteration. From our analysis, IDH1 mutant gliomas are likely to retain a greater degree of methylation compared to IDH wildtype glioma, underscoring a potential link between epitranscriptomic remodeling and glioma pathogenesis.

Our findings are also supported by regional brain m6A profiling studies. A study investigating the isoform level m6A profiling in postmortem human brain regions (prefrontal cortex, caudate nucleus, cerebellum) reported detection of over 57,000 m6A sites, corresponding to 27% of expressed mRNA^16^. Brain region specific distribution of m6A sites has also been examined in gliomas. A recent study by Steponaitis et al. reported a higher overall prevalence of m6A modifications in the cerebellum compared to other brain regions^17^. Finally, there is considerable evidence for high enrichment of m6A in the CNS^18–20^.

Integrated analysis of m6A site distribution and gene expression revealed context-dependent regulatory effects. Since genes comprise transcripts of varying lengths and biotypes, the impact of m6A appears to be modulated by both transcript structure and the region of modification. In the glioma model, gene expression changes were strongly associated with the CDS:3′UTR m6A ratio. Specifically, higher m6A density in the CDS of naive, untreated cells correlated with gene upregulation. In contrast, ALKBH5 KD cells showed a shift in m6A localization toward the 3′UTR, which was associated with increased gene expression. These findings suggest that changes in regional m6A distribution, particularly reduced CDS enrichment, may influence transcript stability and expression dynamics. The m6A redistribution upon regulator KD was evident in both shared and unique m6A sites, with the highest CDS enrichment noted in naive, untreated glioma cells.

Significant isoform switching was observed across all conditions, particularly in astrocytes vs. naive and ALKBH5 KD vs. naive comparisons. Protein-coding isoforms were preferentially used in astrocytes, while retained intron and nonsense mediated decay isoforms were enriched in the KD conditions.

ALKBH5 KD, in particular, was enriched for NMD isoforms, while IGF2BP2 KD and METTL3 KD showed elevated retained intron usage. Structural consequence analysis identified recurrent ORF loss, domain loss, and UTR shortening, many of which have been associated with impaired protein function.

Prior studies have implicated the role of IGF2BP2^21^, METTL3^22^, and ALKBH5^23^ in contributing to alternative splicing. RNA m6A modifications can influence alternative splicing by recruiting specific RNA-binding proteins (RBPs) or modulating their interactions with target RNAs^24,25^. Conversely, splicing events can affect m6A deposition and recognition^24^. Splicing dysregulation has been linked to glioma progression^26^ and treatment resistance^27^.

Our findings expand on this by showing that m6A regulator perturbation not only alters splicing factor expression (e.g., PTBP1, SRSF family) but also shifts isoform usage toward noncoding or decay-prone transcripts. We observed differential expression of a key RBP, PTBP1 (polypyrimidine tract binding protein 1). PTBP1, known to be involved in the regulation of alternative splicing^28,29^, was significantly upregulated in both astrocytes and ALKBH5 KD conditions relative to naive. A study by Yang et al. identified upregulation of PTBP1 in IDH1 wild-type GBM compared with IDH1 mutant gliomas^28^.

Another recent study determined that PTBP1 overexpression acts as a tumor suppressor in glioma by promoting ferroptosis^30^. In addition, members of the SRSF family (SRSF1/2/6) were more highly expressed in astrocytes. Among decay factors, the decapping enzyme 2 (DCP2), a proposed immune related glioma biomarker^31^, was consistently upregulated across all KD conditions. We also noted increased expression of EXOSC10 in astrocytes, ALKBH5 KD, and IGF2BP2 KD cells. EXOSC10 is a core component of the RNA exosome complex. It plays a central role in RNA processing and degradation^32^. Depletion of EXOSC10 has been linked to accumulation of damage-induced long non-coding RNA^32^.

To understand how m6A dysregulation alters broader transcriptomic distribution, we examined the expression of m6A regulators across conditions. In addition to expected changes following knockdowns, we observed global shifts in the expression of other regulators, reflecting coordinated crosstalk among various components of the m6A machinery. Astrocytes showed higher expression of writers (e.g., METTL3, METTL14) and lower expression of erasers (ALKBH5, FTO) relative to glioma cells. IGF2BP2 and the transcription factor ZNF217 were both downregulated in astrocytes and ALKBH5 KD cells. Importantly, IGF2BP2 overexpression is associated with poor prognosis in glioma^33–35^. The transcription factor, ZNF217 mediates tumorigenesis by controlling m6A deposition in embryonic stem cells^36^.

Finally, our data also revealed an additional layer of m6A mediated regulation of gene expression. Beyond transcript biotype and regional distribution, the functional pathway context and cellular state further influence the strength and direction of the correlation between m6A and gene expression.

To elucidate downstream functional consequences, we examined m6A-mediated regulation of gene expression across key pathways. A positive correlation between transcript-level m6A hypermethylation and gene expression was observed in several signaling pathways, including MAPK, mTOR, MYC, and apoptosis. These pathways are central to glioma development, progression, and therapeutic resistance. Our results highlight a dynamic role for m6A in shaping glioma cell behavior by modulating the expression of critical regulatory pathways and suggest that targeted manipulation of m6A deposition or recognition could offer novel therapeutic strategies for selectively disrupting oncogenic signaling.

Our study provides a comprehensive view of m6A mediated transcriptomic and genomic regulation in glioma, integrating gene expression, isoform dynamics, and functional pathway analysis in the context of targeted knockdown of m6A machinery. We report distinct patterns of m6A deposition, regional redistribution, and their downstream effects on RNA stability, splicing, and decay. Importantly, these changes were not uniform but highly context-dependent, influenced by transcript biotype, modification location, and cellular state. Our findings advance the understanding of how epitranscriptomic remodeling contributes to glioma pathogenesis and progression. Future studies are needed to study the potential of targeting m6A mediated regulatory networks as a therapeutic strategy in glioma and other malignancies.

## Methods

### Study Population

Human glioma cell line, Gli36 (generated at Massachusetts General Hospital with approved IRB procedures) was utilized for in vivo m6A RNA methylome profiling. Knockdown (KD) of three distinct m6A regulators was achieved by silencer RNA (siRNA) mediated transfection. A total of n = 5 experimental conditions were tested: (i) naive, no treatment, (ii) ALKBH5 (eraser) KD, (iii) IGF2BP2 (reader) KD, (iv) METTL3 (eraser) KD, and (v) healthy, nondiseased astrocytes. Three technical replicates were performed per condition, yielding a total of N = 15 sequencing runs.

### Selected Silencer RNA (siRNA) for m6A regulator knockdown

Select Pre-designed siRNAs (Catalog Number: 4427037) were purchased from ThermoFisher Scientific for knockdown of known m6A regulators: ALKBH5 (siRNA ID: s29688), IGF2BP2 (siRNA ID: s20924), and METTL3 (siRNA ID: s32142).

### Cell line transfection and collection

On Day 0 (one day before transfection), 2 x 105 Gli36EGFRvIII cells were plated in a 6-well plate using Opti-MEMTM reduced serum medium (ThermoFisher Scientific, Waltham, MA, USA). The following day, cells were checked to ensure 60-80% confluency prior to transfection. Transfection on Day 1 was performed using Lipofectamine RNAiMAX Reagent (ThermoFisher Scientific, Waltham, MA, USA) according to manufacturer’s instructions. Briefly, both lipofectamine RNAiMAX reagent and silencer RNA (siRNA) concentrates were diluted in Opti-MEM medium. A volume of 3 μl of 10 μM siRNA was combined with 150 μl of Opti-MEM medium to achieve a final concentration of 30 pmol. RNA-lipid complexes were then prepared by combining diluted siRNA and diluted lipofectamine RNAiMAX reagent in a 1:1 ratio followed by 5 min incubation at room temperature. The siRNA-lipid complex was then added to the individual wells of the 6 well plate. The reaction mixture was mixed gently by rocking the plate back and forth. Cells were transfected for 48h at 37°C in a CO2 incubator to achieve optimal m6A regulator knockdown. On Day 3, the Opti-MEM medium was carefully aspirated and replaced with Dulbecco’s Modified Eagle Medium (DMEM, Invitrogen, Waltham, MA) with high glucose containing 10% fetal bovine serum (FBS; Life Technologies Corporation, Carlsbad, CA, USA) and 1% Penicillin/Streptomycin solution (Pen/Strep; Life Technologies Corporation, Carlsbad, CA, USA).

Transfected cells were cultured in DMEM for 24h and transferred to T-75 flask on Day 4. On Day 7, cells were transferred to Nunc™ TripleFlask™ Treated Cell Culture Flasks for cell culture expansion. Cell viability and number was assessed using the Countess II FL Automated Cell Counter (ThermoFisher Scientific, Waltham, MA, USA). For downstream analytics, cells were pelleted to create multiple aliquots (3 x 106 cells per aliquot) and stored at −80°C until further analysis.

### Mycoplasma Contamination testing

Testing for mycoplasma contamination was performed using a commercial mycoplasma PCR kit (PCR Mycoplasma Detection kit; Applied Biological Materials Incorporated, BC, Canada). The testing was done at two stages of the workflow: (i) Day 3, end of 48h transfection period, and (ii) Day 7, prior to transfer of cells from T-75 flask to Nunc™ TripleFlask™.

### Assessment of transfection efficiency

Quantification (percentage gene knockdown) and stability of transfection were assessed using quantitative polymerase chain reaction (qPCR) measurement of the target. To ensure stability of transfection, quantification was performed using cells collected from three stages of the workflow: (i) 6 well plate, (ii) T-75 flask, and (iii) Nunc™ TripleFlask™. Commercially available, pre-designed TaqMan gene expression assays were purchased (ThermoFisher Scientific, Waltham, MA, USA) for ALKBH5, IGF2BP2, and METTL3. Isolated total RNA from transfected (KD, control) cells was reverse transcribed using SuperScript™ VILO™ cDNA Synthesis Kit (ThermoFisher Scientific, Waltham, MA, USA). The qPCR reaction was performed in the Applied Biosystems StepOnePlus Real-Time PCR (Thermo Fisher Scientific) in a final volume of 20 μl and was prepared using 10 μl of TaqMan Universal Master Mix II, no UNG (Thermo Fisher Scientific), 1 μl of 20X gene expression assay, and cDNA. Each sample was run in three replicates. The data was analyzed by visualizing amplification plots and comparing the Cycle threshold (Ct) values in knockdown (ALKBH5, IGF2BP2, METTL3) and control (scrambled siRNA) wells. Normalization was performed using GAPDH as a reference assay (ThermoFisher Scientific).

### Cellular RNA Extraction and Quantification

Total RNA was extracted from transfected cells using TRIzol lysis reagent (ThermoFisher Scientific, Waltham, MA, USA) per manufacturer’s protocol. For each condition, 3 x 106 million cells were lysed using 3 ml TRIzol. The extracted RNA was then eluted in xx μl of nuclease free water (Qiagen). Agilent RNA 6000 Pico Kit was used with Agilent Technologies 2100 Bioanalyzer to determine the concentration and RIN (RNA Integrity Number) value of the samples. Isolated RNA was stored at −80°C prior to downstream analysis.

### Cellular and EV messenger RNA (mRNA) enrichment

Following Cell RNA extraction, total RNA was selectively enriched for mRNA using RiboMinus™ Eukaryote Kit v2 (ThermoFisher Scientific, Waltham, MA, USA) per manufacturer’s recommendations. This provides transcriptome isolation downstream sequencing by selectively depleting 5S, 5.8S, 18S, and 28S ribosomal RNA from total RNA. For high yield cell RNA, RiboMinus™ Eukaryote Kit v2 (Catalog ID: A15026) was used to maximize the mRNA enrichment and overall recovery.

### RNA Quality Control Steps

Different measures were integrated throughout the workflow for quantitative and qualitative analysis of cell RNA and EV RNA. Quantification of “isolated total RNA” and “mRNA post rRNA depletion” was performed using Bioanalyzer Agilent RNA 6000 Pico Kit. Overnight ethanol precipitation with 3M sodium acetate at −20°C was performed post RNA extraction and post rRNA depletion to prevent carry over inhibitors during library preparation.

### Library Preparation and Sequencing

Direct RNA sequencing of Cell derived mRNA was performed using the Oxford Nanopore technologies (ONT) platforms. Libraries were prepared using SQK-RNA002 (ONT) kit per manufacturer’s protocol. The library was then sequenced on the MinION software using flow cells 9.4.1 (FLO-MIN106D, ONT). No multiplexing was done and each sample was run individually for (24-72)h.

## Data Analysis

### Preprocessing

Raw RNA sequencing reads from cells and extracellular vesicles (EVs), generated using Oxford Nanopore Technologies (ONT), were basecalled from POD5 files using Dorado (v0.5) with default parameters. The decoder file “rna002_70bps_fast@v3” was employed to convert raw signal data into FASTQ format, optimized for RNA sequencing at 70 base pairs per second. Reads that passed Dorado’s default quality filters were retained for downstream analysis. Due to a later sequencing batch, astrocyte cell samples were basecalled using the updated decoder “rna004_130bps_fast@v3.0.1” to ensure compatibility with newer ONT flow cells. The basecalled reads were then aligned to the human transcriptome (GENCODE v45, GRCh38) using minimap2 (v2.17). All passing aligned reads were included in the analysis. The BAM files were sorted and indexed using Samtools (v1.10). The processed and indexed BAM files were then used for transcript quantification and subsequent analyses. Per condition, plots were computed to compare average read length vs. read quality (Supplementary Figure 1). Non weighted histograms of read lengths after log transformation were also compared across all experimental conditions (Supplementary Figure 2).

### Quantification

Transcript and gene quantification was performed using bambu (v3.11.1), which supports the quantification of both known and novel transcripts and genes, as well as full length transcripts from long-read RNA sequencing data. BAM files generated by minimap2 were used as input along with the GENCODE v45 GRCh38 genome and annotation files. Transcript expression levels were quantified for each sample, with both raw counts and counts per million (CPM) values calculated. Bambu was configured to detect novel transcript isoforms in addition to quantifying known transcripts. To ensure robust quantification, transcripts with zero counts were excluded from analysis in the Cell RNA data, and transcripts with zero counts in more than two samples across all experimental conditions were excluded from the analysis in the EV RNA data. This filtering step reduced the noise and ensured that low-abundance transcripts did not affect downstream analyses.

### Differential Expression Analysis

Using DESeq2 (v1.48.1), differential gene expression (DGE) analysis was performed on the transcripts and gene counts generated by bambu. To account for biological differences and avoid confounding effects, DGE analysis was conducted in two separate groups. First, to examine the broad transcriptomic differences between cell types, astrocytoma cells were compared to naive glioma cells (no knockdown). Second, to assess the effects of individual m6A regulator knockdowns, a separate DESeq2 analysis was performed comparing naive glioma cells and each knockdown condition (ALKBH5 KD, IGF2BP2 KD, METTL3 KD). This comparison excluded the astrocyte samples, as preliminary analyses indicated that a combined design matrix introduced significant batch effects and minimized the DGE signal among the knockdown and naive conditions. By excluding astrocyte samples in the knockdown analysis, we maximized sensitivity for detecting knockdown-specific transcriptomic changes. For each comparison, genes and transcripts with less than 1 count across all samples were filtered out, and remaining counts were normalized using DESeq’s internal size factor estimation. Transcripts and genes with a fold change of at least 1.5 (log2fFC > 0.58, log2FC < −0.58) and an adjusted p-value (Benjamini-Hochberg) less than 0.05 were considered significantly differentially expressed. EnhancedVolcano (v1.24) in R was used to visualize fold changes and statistical significance for each pairwise comparison. pheatmap (v1.0.12) was used to generate heatmaps of log2 fold changes for genes and transcripts corresponding to (i) known m6A regulators, (ii) RNA splicing, and (iii) RNA decay. For RNA splicing and RNA decay, unsupervised hierarchical clustering using Euclidean distance was applied to rows to identify shared expression profiles across conditions. Significant log2 fold changes (p-value < 0.05) were denoted with asterisks.

### Isoform usage analysis

Differential isoform usage analysis was performed in R using IsoformSwitchAnalyzeR38 (2.8.0). The aggregated isoform counts and abundances were input along with the human (GRCh38.p14) annotation and transcriptome files. Single isoform genes were filtered out during the preFilter() step since these genes cannot exhibit changes in isoform usage. Statistical analysis was performed with isoformSwitchTestDEXSeq(). We required a difference in isoform proportions between classification groups of >0.1 and an FDR-adjusted p-value of <0.05 for significance. The functional consequences of the identified isoform switches were generated using the analyzeSwitchConsequences() function. This analysis identified changes in key functional domains such as coding potential, exon loss, domain loss, and isoform length where the difference in isoform proportions is greater than 0.4. Alternative splicing analysis was performed using the analyzeAlternativeSplicing() function, which identified and assessed the significance of specific events such as alternative 3’ acceptor site (A3) gain or loss. Consequence plots were generated that show the fraction of significant isoform switches that have a specific consequence and its opposing consequence in each comparison.

### Transcriptome-wide m6A-modified sites

RNA methylation analysis was conducted using m6anet (v2.1.0), a machine learning-based tool, to detect RNA modifications at single-nucleotide resolution. BAM files obtained from minimap2 were input to m6anet, along with the raw FAST5 files, FASTQ, and transcriptome files. The model was trained using the RNA004 direct RNA-Seq run using HEK293T cell line. As m6anet supports pooling over replicates, the replicates (n=3) per condition were pooled prior to the inference step. The output of m6anet predicted m6A sites with probability that the site is modified, the transcript position of the site, the 5-mer motif of the site, and the estimated percentage of reads in the site that is modified. Sites with a probability of modification greater than 0.9 were considered high confidence m6a sites and retained for downstream analysis. Post processing of m6a site predictions was conducted using a custom R package, m6ASeqTools, to standardize and streamline downstream processes. Key metrics such as the distribution of modified sites, transcripts, and genes, the frequency of transcript biotypes, k-mer enrichment, and the length distribution of modified transcripts were computed and visualized.

### Distribution of modified sites

m6ASeqTools was used to calculate the lengths of the 5’ UTR, CDS, and 3’ UTR regions in order to analyze the distribution of m6A-modified sites across transcript regions. For each transcript, the coding sequence (CDS) length was determined by summing exon lengths, while the UTR lengths were calculated based on start and end coordinates. Transcripts with missing UTR annotations were assigned a length of zero for the missing regions. To visualize the distribution of m6A-modified sites across these regions, the transcript position of each modification site was compared with the UTR and CDS regions to determine if the site was located within the 5’ UTR, CDS, or 3’ UTR. The relative position within each region was calculated, and kernel density estimation (KDE) plots were generated to visualize the relative density of m6A sites across these transcript regions. Modifications grouped by transcript biotypes (protein-coding, nonsense mediated decay, retained intron) and methylation status were visualized.

### Identification of hyper- and hypo-m6A methylated Transcripts

To identify hyper- and hypo-m6A methylated transcripts, we first filtered for transcripts present in all knockdown conditions (ALKBH5, IGF2BP2, and METTL3) vs. Naive group. Then, we filtered for transcripts present in the Astrocytes vs. Naive group. For these commonly identified sites, we then identified transcripts containing sites common to all comparisons. For these commonly modified transcripts, we calculated the average modification ratio for each site within each classification. Finally, to calculate the weighted modification ratio for these commonly modified transcripts, we summed the modification ratios of all sites within each transcript and divided by the transcript length. This approach ensured that transcripts with more modified sites contributed proportionally while preventing length bias, allowing for a more accurate comparison of modification levels across transcripts of varying lengths. To identify hypermethylated and hypomethylated transcripts, we calculated the log2 fold change of the weighted modification ratio for each pairwise comparison (n=7). A positive log2FC indicated that a transcript was hypermethylated in Naive (vs. Astrocytes), ALK (vs. METTL3, IGF2BP2, Naive), IGF2BP2 (vs. METTLE3, Naive), or METTL3 (vs. Naive), while a negative log2FC indicated that a transcript was hypermethylated in Astrocytes (vs. Naive), METTLE3 (vs. ALKBH5, IGF2BP2), Naive (vs. ALKBH5, IGF2BP2, METTLE3), or IGF2BP2 (vs. ALKBH5). ggplot2 (v3.5.2) in R was used to illustrate the different biotypes distribution in each cluster by plotting a stacked barchart for all pairwise comparisons. For each comparison, a line graph was generated to show the relationship between the ratio of the abundance in the CDS region compared to the 3’UTR region where there is significant methylation.

### Functional Analysis of Isoform Switching and hyper- and hypo-m6A methylated Genes

Functional analysis was performed using ClusterProfiler39 (4.14.6) for all Gene ontologies (Biological Process, Molecular Function, and Cellular Component) and KEGG pathways. The enrichGO function was applied for genes that were upregulated and had m6A modifications in each tumor classification. For each comparison, plots were generated that show the amount of differentially expressed genes associated with the top twenty pathways based on fold enrichment score. Heatmaps of differential gene expression and the associated m6a modifications were generated for selected enriched pathways (i.e. Apoptosis, MAPK Signaling, Oxidative Phosphorylation).

### Gene Set Enrichment Analysis

Gene Set Enrichment Analysis was used to show pathways that are significantly associated within the selected gene list. Gene set enrichment analysis is used to determine if an inputted gene list is overly expressed in the enriched pathway.

### Institutional review board statement

Our studies were conducted in accordance with principles for human experimentation as defined in the U.S. Common Rule and were approved by the Human Investigational Review Board of each study center under Partners institutional review board (IRB)-approved protocol number 2017P001581. All healthy control subjects were screened for pertinent oncologic and neurologic medical histories. Individuals with a history of cancer, neurological disorders, and infectious diseases were excluded from the study.

### Data availability

The sequencing and m6A modification data generated in this study are provided in the Supplementary Information and Source Data file. The raw nanopore sequencing data and m6Anet output are not publicly available but will be available upon request from the corresponding author. Source data are provided with this paper.

## Acknowledgements

The authors would like to thank all the MGH Neurosurgery clinicians and staff who assisted with the collection of samples. We are also deeply appreciative to the patients and their families for participating in the study. This work is supported by grants R01 CA239078, CA237500, CA291826 (LB). MGN Transformative Scholar (LB), Rappaport Scholar (LB). The funding sources had no role in the writing of the manuscript or the decision to submit the manuscript for publication. The authors have not been paid to write this article by any entity. The corresponding author has full access and assumes final responsibility for the decision to submit for publication.

## Contributions

S.M.B., H.L., A.K.E, S.M.K. L.B., Data curation, formal analysis, validation, investigation, visualization, writing of original draft, writing, review and editing. A.K.E., T.H., Experimental work, analysis, review and editing; D.G., K.F., A.K., Sample collection, processing, clinical correlates, data analysis: B.S.C., Formal analysis, methodology, review and editing. L.B., Conceptualization, resources, data curation, software, formal analysis, supervision, funding acquisition, validation, investigation, visualization, methodology, writing—original draft, project administration, writing—review and editing.

## Corresponding author

Correspondence to Leonora Balaj.

## References

1. Jiang, X. et al. The role of m6A modification in the biological functions and diseases. Signal Transduct Target Ther 6, 74 (2021).

2. Akhtar, J., Lugoboni, M. & Junion, G. mA RNA modification in transcription regulation. Transcription 12, 266–276 (2021).

3. Huang, W. et al. N6-methyladenosine methyltransferases: functions, regulation, and clinical potential. J. Hematol. Oncol. 14, 117 (2021).

4. Jiang, L., Li, X., Wang, S., Yuan, Z. & Cheng, J. The role and regulatory mechanism of mA methylation in the nervous system. Front. Genet. 13, 962774 (2022).

5. Fan, Y., Lv, X., Chen, Z., Peng, Y. & Zhang, M. m6A methylation: Critical roles in aging and neurological diseases. Front Mol Neurosci 16, 1102147 (2023).

6. Louis, D. N. et al. The 2021 WHO Classification of Tumors of the Central Nervous System: a summary. Neuro Oncol 23, 1231–1251 (2021).

7. Xu, S., Tang, L., Li, X., Fan, F. & Liu, Z. Immunotherapy for glioma: Current management and future application. Cancer Lett 476, 1–12 (2020).

8. Mohammed, S., Dinesan, M. & Ajayakumar, T. Survival and quality of life analysis in glioblastoma multiforme with adjuvant chemoradiotherapy: a retrospective study. Rep Pract Oncol Radiother 27, 1026–1036 (2022).

9. Chen, J.-J. et al. The m6A reader HNRNPC promotes glioma progression by enhancing the stability of IRAK1 mRNA through the MAPK pathway. Cell Death Dis 15, 390 (2024).

10. Zhang, Y. et al. m6A modification in RNA: biogenesis, functions and roles in gliomas. J Exp Clin Cancer Res 39, 192 (2020).

11. Suvà, M. L. et al. Reconstructing and reprogramming the tumor-propagating potential of glioblastoma stem-like cells. Cell 157, 580–594 (2014).

12. Yuan, F. et al. Roles of the mA Modification of RNA in the Glioblastoma Microenvironment as Revealed by Single-Cell Analyses. Front Immunol 13, 798583 (2022).

13. Cui, Q. et al. mA RNA Methylation Regulates the Self-Renewal and Tumorigenesis of Glioblastoma Stem Cells. Cell Rep 18, 2622–2634 (2017).

14. Vitting-Seerup, K. & Sandelin, A. IsoformSwitchAnalyzeR: analysis of changes in genome-wide patterns of alternative splicing and its functional consequences. Bioinformatics 35, 4469–4471 (2019).

15. Batool, S. M. et al. Epitranscriptome mapping of m^6^A RNA modifications in glioma tumor tissue. medRxiv (2024) doi:10.1101/2024.09.24.24314089.

16. Gleeson, J., et al. Isoform-level profiling of m6A epitranscriptomic signatures in human brain. bioRxiv (2024) doi:10.1101/2024.01.31.578088.

17. Steponaitis, G. et al. m6A-lncRNA landscape highlights reduced levels of m6A modification in glioblastoma as compared to low-grade glioma. Mol Med 31, 195 (2025).

18. Du, B. et al. N6-methyladenosine (m6A) modification and its clinical relevance in cognitive dysfunctions. Aging (Albany NY) 13, 20716–20737 (2021).

19. Chokkalla, A. K., Mehta, S. L. & Vemuganti, R. Epitranscriptomic regulation by mA RNA methylation in brain development and diseases. J Cereb Blood Flow Metab 40, 2331–2349 (2020).

20. Yu, J., She, Y. & Ji, S.-J. mA Modification in Mammalian Nervous System Development, Functions, Disorders, and Injuries. Front Cell Dev Biol 9, 679662 (2021).

21. Liu, W. et al. IGF2BP2 orchestrates global expression and alternative splicing profiles associated with glioblastoma development in U251 cells. Transl Oncol 51, 102177 (2025).

22. Kim, Y. et al. METTL3 regulates alternative splicing of cell cycle-related genes via crosstalk between mRNA mA modifications and splicing factors. Am J Cancer Res 13, 1443–1456 (2023).

23. Tang, C. et al. ALKBH5-dependent m6A demethylation controls splicing and stability of long 3’-UTR mRNAs in male germ cells. Proc Natl Acad Sci U S A 115, E325–E333 (2018).

24. Zhu, Z.-M., Huo, F.-C., Zhang, J., Shan, H.-J. & Pei, D.-S. Crosstalk between m6A modification and alternative splicing during cancer progression. Clin Transl Med 13, e1460 (2023).

25. Mehravar, M. & Wong, J. J.-L. Interplay between N-adenosine RNA methylation and mRNA splicing. Curr Opin Genet Dev 87, 102211 (2024).

26. Song, X. et al. RNA splicing analysis deciphers developmental hierarchies and reveals therapeutic targets in adult glioma. J Clin Invest 134, (2024).

27. Ben Mrid, R., El Guendouzi, S., Mineo, M. & El Fatimy, R. The emerging roles of aberrant alternative splicing in glioma. Cell Death Discov 11, 50 (2025).

28. Yang, J. et al. PTBP1-mediated repression of neuron-specific CDC42 splicing constitutes a genomic alteration-independent, developmentally conserved vulnerability in IDH-wildtype glioblastoma. Funct Integr Genomics 24, 135 (2024).

29. Ye, Z., Zhong, Y. & Zhang, Z. Pan-cancer multi-omics analysis of PTBP1 reveals it as an inflammatory, progressive and prognostic marker in glioma. Sci Rep 14, 14584 (2024).

30. Sun, J. et al. PTBP1 acts as a tumor suppressor in glioma by promoting HMOX1-dependent ferroptosis. Biochem Pharmacol 239, 117041 (2025).

31. Mei, Y. et al. Decapping enzyme 2 is a novel immune-related biomarker that predicts poor prognosis in glioma. Biotechnol Genet Eng Rev 40, 4262–4283 (2024).

32. Domingo-Prim, J. et al. EXOSC10 is required for RPA assembly and controlled DNA end resection at DNA double-strand breaks. Nat Commun 10, 2135 (2019).

33. Li, N. et al. IGF2BP2 modulates autophagy and serves as a prognostic marker in glioma. Ibrain 10, 19–33 (2024).

34. Zhou, H.-L., Chen, D.-D. & Li, X.-L. Pan-cancer analysis identified IGF2BP2 as a potential prognostic biomarker for multiple tumor types. Egypt. J. Med. Hum. Genet. 25, (2024).

35. Liu, H. et al. mA reader IGF2BP2-stabilized CASC9 accelerates glioblastoma aerobic glycolysis by enhancing HK2 mRNA stability. Cell Death Discov 7, 292 (2021).

36. Lee, D.-F., Walsh, M. J. & Aguiló, F. ZNF217/ZFP217 Meets Chromatin and RNA. Trends Biochem Sci 41, 986–988 (2016).

