## Supplemental data for "Deregulating m6A regulators leads to altered RNA biology in glioma cell lines"

#### Supplementary Figure 1

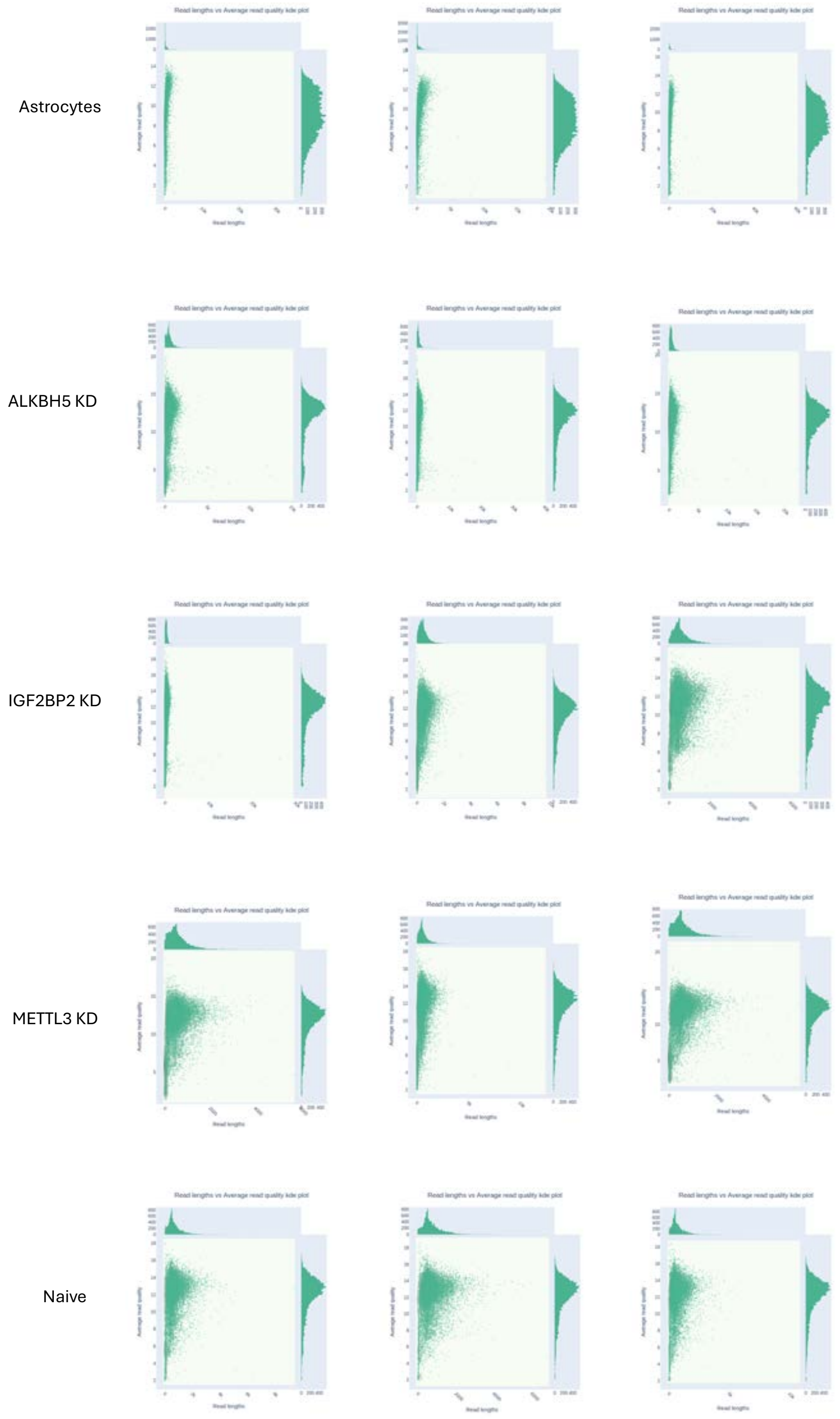


**Supplementary Figure 1: Sequencing Read Quality and Length Analysis**
Quality control (QC) Nanoplots plotting the log transformed read lengths against average read quality (using a kernel density estimate) in Astrocytes (n=3), ALKBH5 KD (n=3), IGF2BP2 KD (n=3), METTL3 KD (n=3), and Naïve (n=3) conditions.

**Supplementary Figure 2**

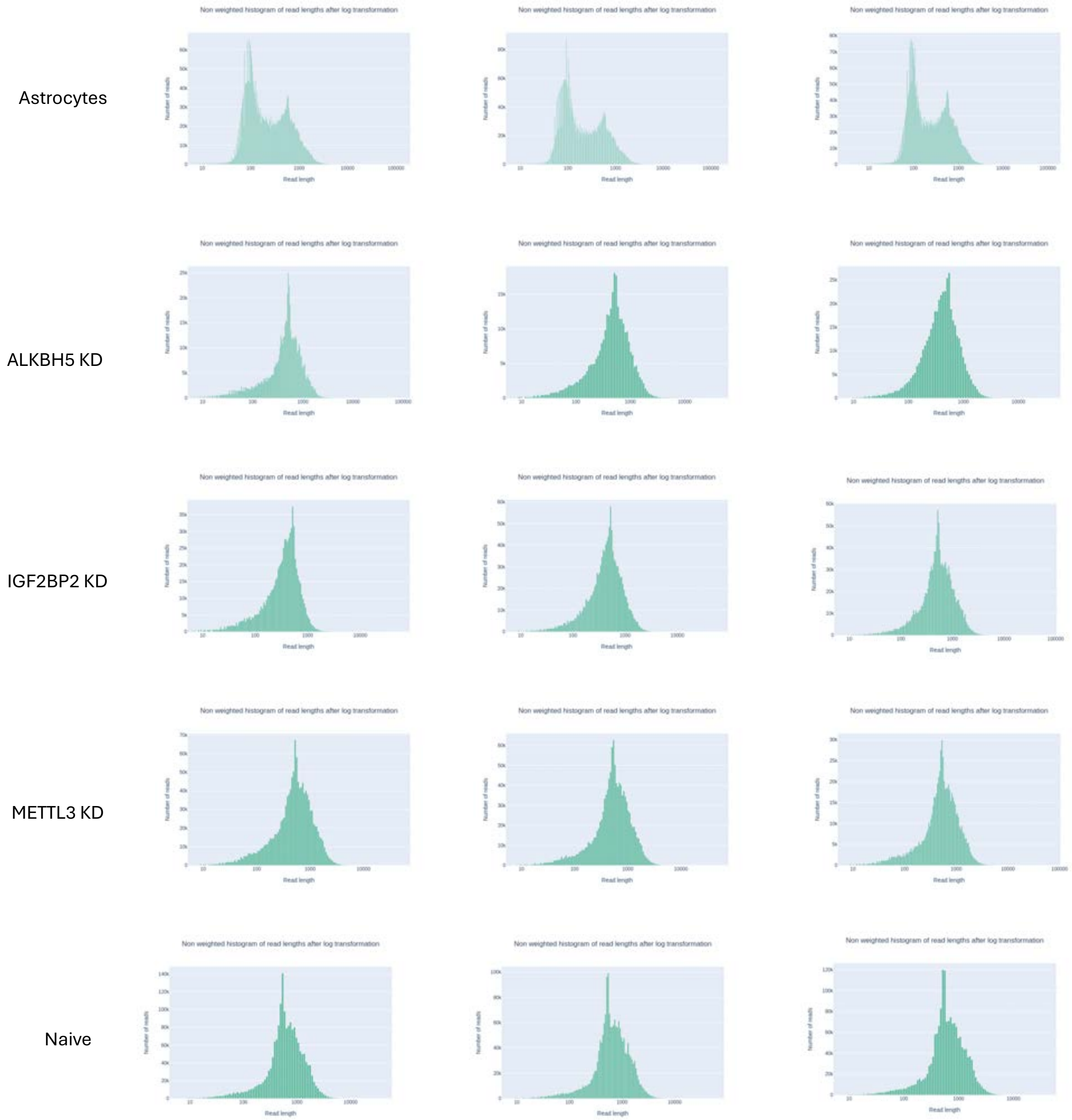


**Supplementary Figure 2: Distribution of Read Lengths.**
Distribution of log transformed read lengths in Astrocytes (n=3), ALKBH5 KD (n=3), IGF2BP2 KD (n=3), METTL3 KD (n=3), and Naïve (n=3) conditions.

#### Supplementary Figure 3

a.

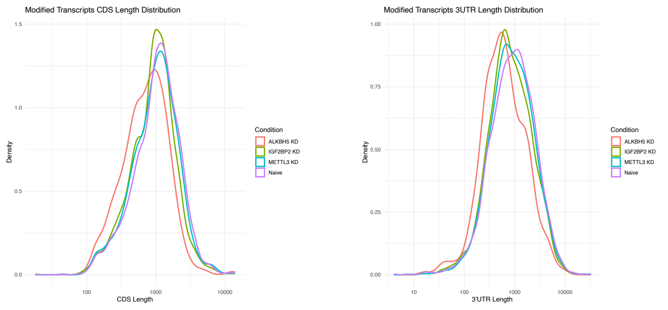


b.

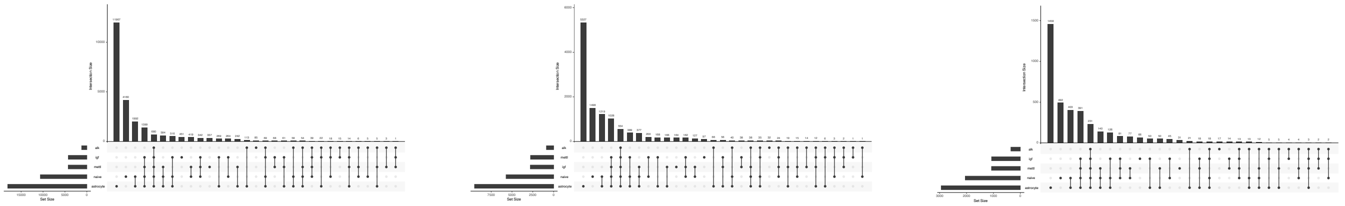

c.

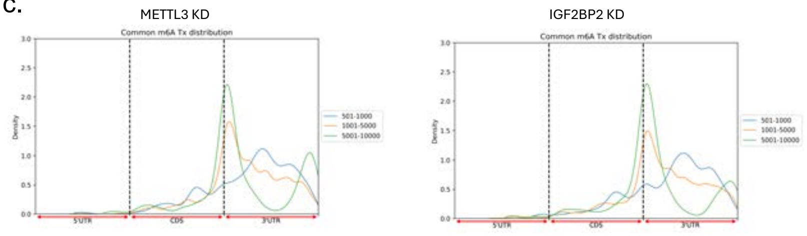


**Supplementary Figure 3:** (a) Distribution of CDS (left) and 3'UTR (right) lengths of the m6A modified transcripts across the 4 conditions. (b) UpSet plot showing overlap of m6A modified sites (left), transcripts (middle), and genes (right) across Naive, IGF2BP2 KD, METTL3 KD, ALKBH5 KD, and Astrocyte conditions. Horizontal bars indicate the total number of m6A sites, transcripts, and genes in each condition. Vertical bars (length) represent the number of sites, transcripts, and genes shared between the selected combinations of conditions shown below. (c) Density plots of m6A distribution along the transcript region, stratified by RNA length.

#### Supplementary Figure 4

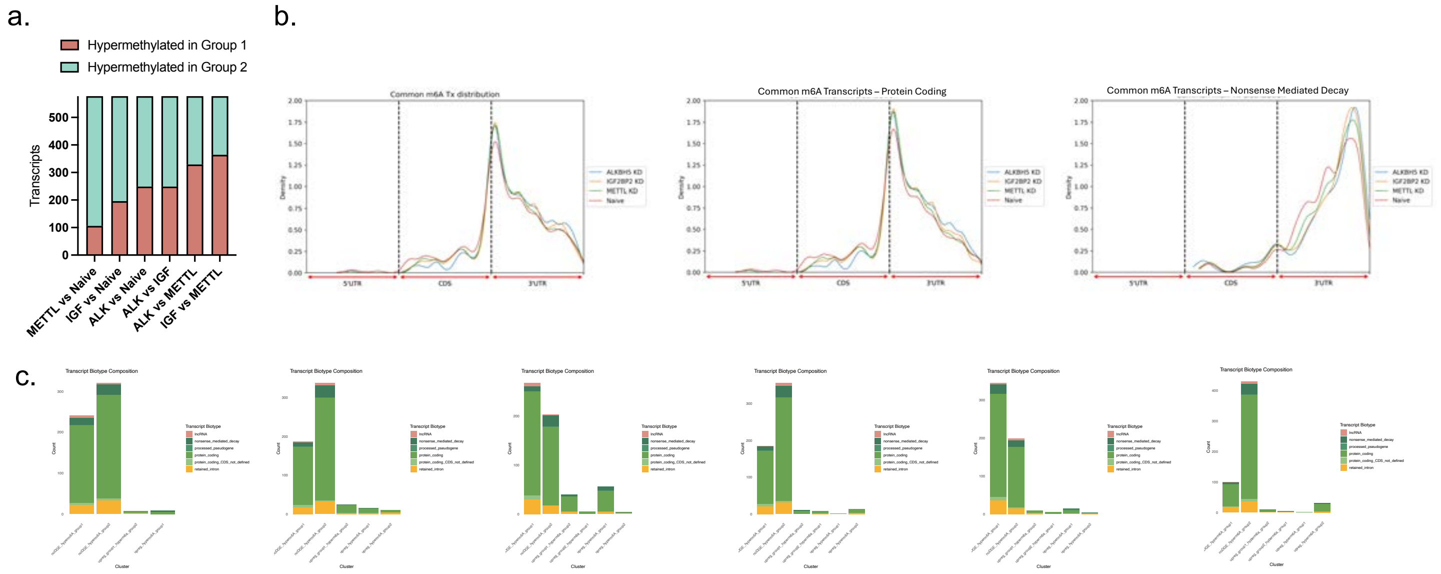


d.

| Commonly modified transcripts (hypermethylated) | | | | |
|---|---|---|---|---|
| Comparison | Group | Gene Upregulated | No change | Gene Downregulated |
| ALK vs IGF | ALK | CDS | 3'UTR | 3'UTR |
| | IGF | NA | 3'UTR | 3'UTR |
| ALK vs METTL | ALK | 3'UTR | 3'UTR | 3'UTR |
| | METTL | 3'UTR | 3'UTR | 3'UTR |
| IGF vs METTL | IGF | CDS | 3'UTR | 3'UTR |
| | METTL | 3'UTR | 3'UTR | 3'UTR |
| ALK vs Naïve | ALK | CDS | 3'UTR | NA |
| | Naïve | CDS | 3'UTR | CDS |
| IGF vs Naïve | IGF | CDS | 3'UTR | 3'UTR |
| | Naïve | 3'UTR | 3'UTR | 3'UTR |
| METTL vs Naïve | METTL | 3'UTR | 3'UTR | 3'UTR |
| | Naïve | 3'UTR | 3'UTR | 3'UTR |

e.

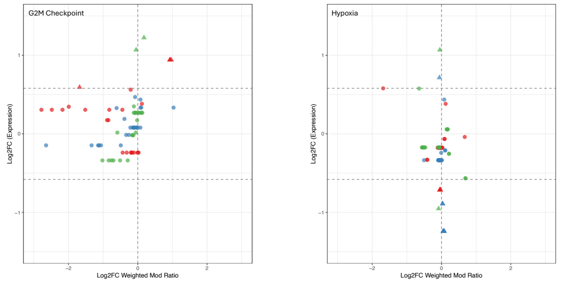


**Supplementary Figure 4:**
(a) Stacked bar graph showing the number of hyper- and hypo-methylated transcripts (n=577) within each comparison (n=6). (b) m6A localization in the 5'UTR, CDS, and 3'UTR for all commonly methylated transcripts within each condition. The middle panel shows the regional distribution for protein-coding transcripts, and the right panel shows the distribution for nonsense-mediated decay transcripts. (c) Biotype distribution for different clusters of upregulated and hypermethylated gene groups per group in comparisons across KD and Naïve conditions. (d) Table displaying only the hypermethylated transcripts in each condition per comparison and whether the associated sites are location more in CDS or 3’UTR depending on whether the gene is upregulated, no change, or downregulated. (e) Scatter plots of selected pathways containing commonly m6A modified genes and displaying differential methylation and gene expression levels between in each knockdown condition compared to naive.

### Supplementary Figure 5

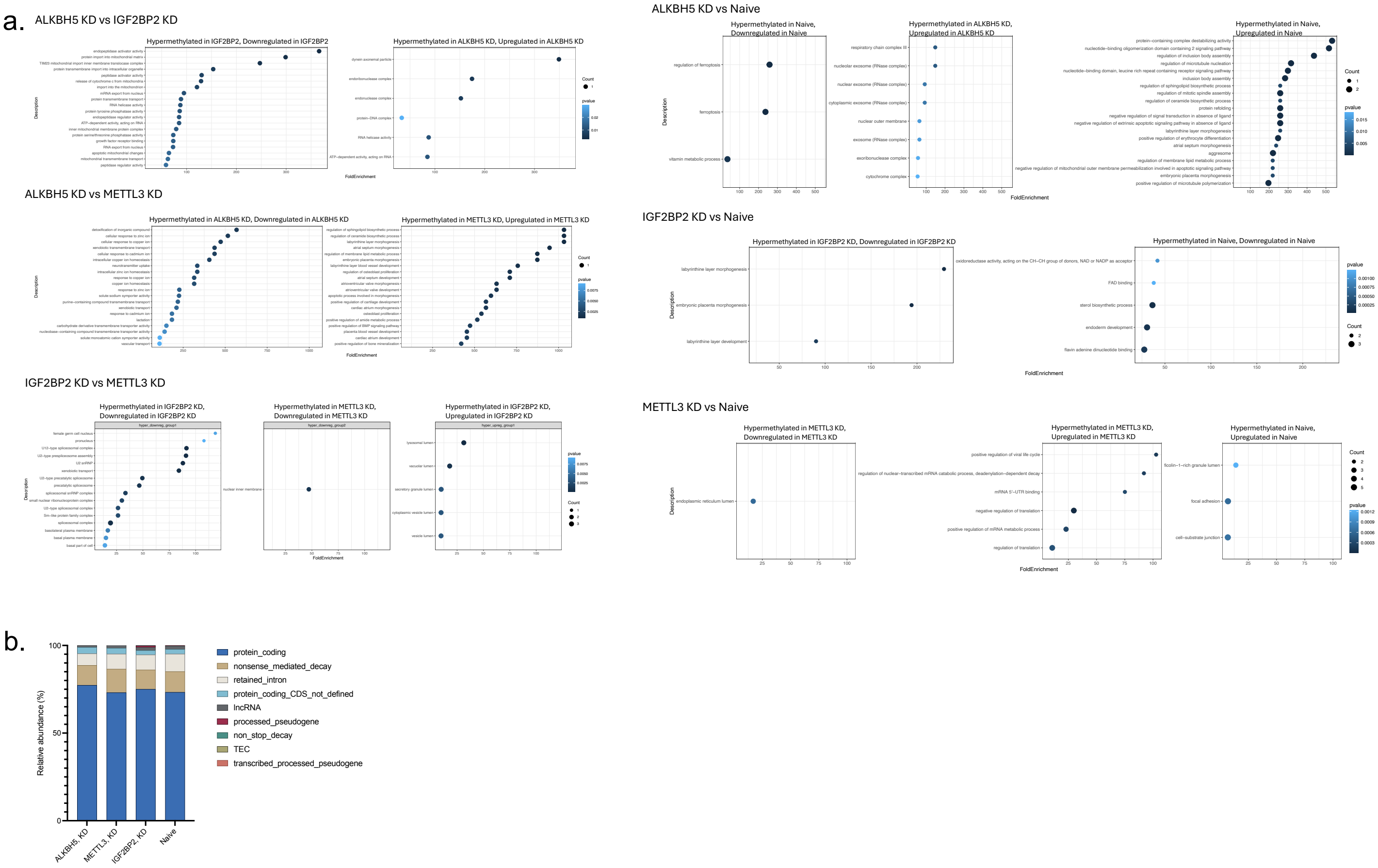


**Supplementary Figure 5: Gene ontology dot plots for the clusters of m6A methylation and gene expression.** (a) Each plot illustrates the gene ontology for specific m6A methylation and gene expression groups with the top 20 pathways based on the fold enrichment score. (b) Biotype distribution of unique m6A sites detected in each KD conditions and controls (naïve, astrocytes).

`

#### Supplementary Figure 6

a.

| Comparison | nrIsoforms | nrSwitches | nrGenes |
|---|---|---|---|
| ALK vs IGF | 6 | 7 | 5 |
| ALK vs METTL | 165 | 172 | 110 |
| ALK vs NAIVE | 1364 | 1253 | 845 |
| IGF vs METTL | 79 | 74 | 53 |
| IGF vs NAIVE | 821 | 777 | 548 |
| METTL vs NAIVE | 459 | 401 | 302 |
| Combined | 2015 | 1903 | 1219 |

c.

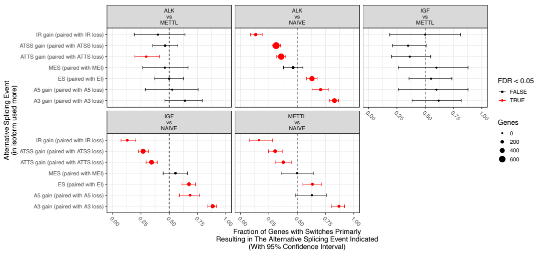


b.

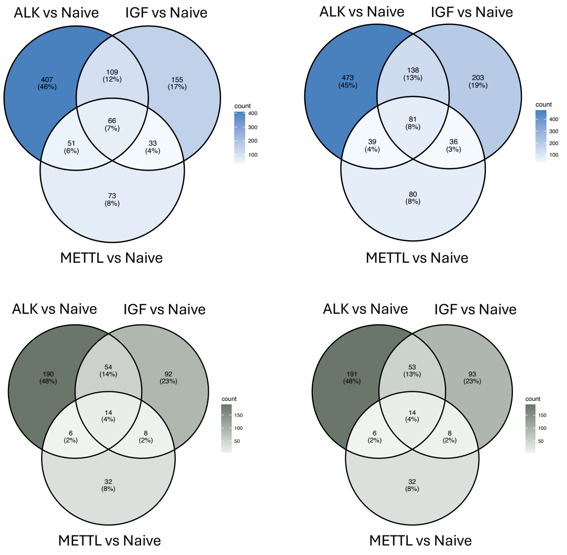


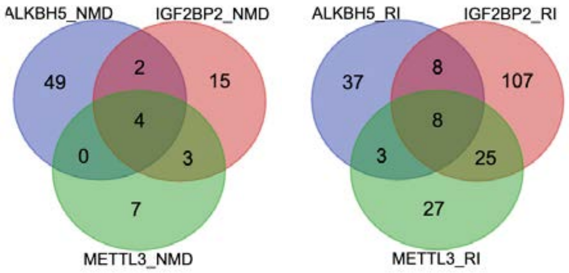


**Supplementary Figure 6: Summary of Isoform Switching for the knockdown comparisons.** (a) Quantification of the isoform switching events to show the number of significant isoforms, switches, and genes for the knockdown comparisons and naïve condition. (b) Venn diagrams of the common and unique significant isoforms and isoforms with a significant consequence divided based on what condition the isoform is used more in. The diagrams on the left side show the isoforms that are used more in the naïve condition and the right show isoforms used more in the knockdown condition. The gray diagrams represent isoforms with a significant consequence, while blue diagrams are a representation of the total significant isoforms. (c) Summary of alternative splicing events with the x-axis showing the ratio of genes with an isoform switch resulting in the alternative splicing event. (d) Summary of isoform switch consequences where values greater than 0.5 indicate that the  and less than 0.5 means the isoforms undergo the opposing consequence more.

**Supplementary Figure 7**

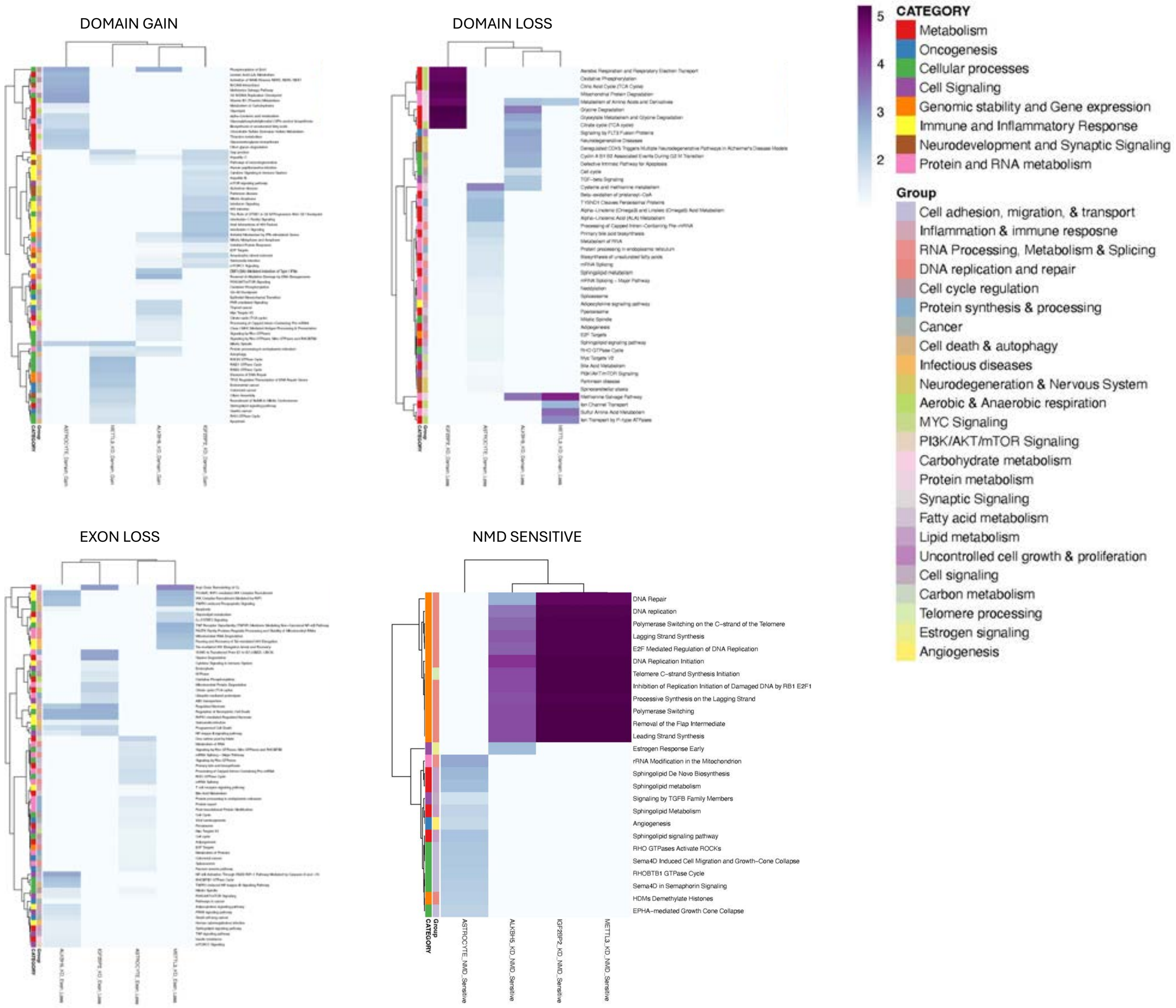


**Supplementary Figure 7: Gene ontology analysis on isoforms of the knockdown and astrocytes condition vs. naïve for a specific consequence of interest.** Four heat maps which show isoforms that are grouped together based on their consequence and a gene ontology study to analyze if there are common or unique pathways that are affected by the same consequence. These pathways are broken down into two groups to give a better overview of how each consequence affects certain pathways. The pathways are ranked based on the log value of the combined score performed by enrichR.

#### Supplementary Figure 8

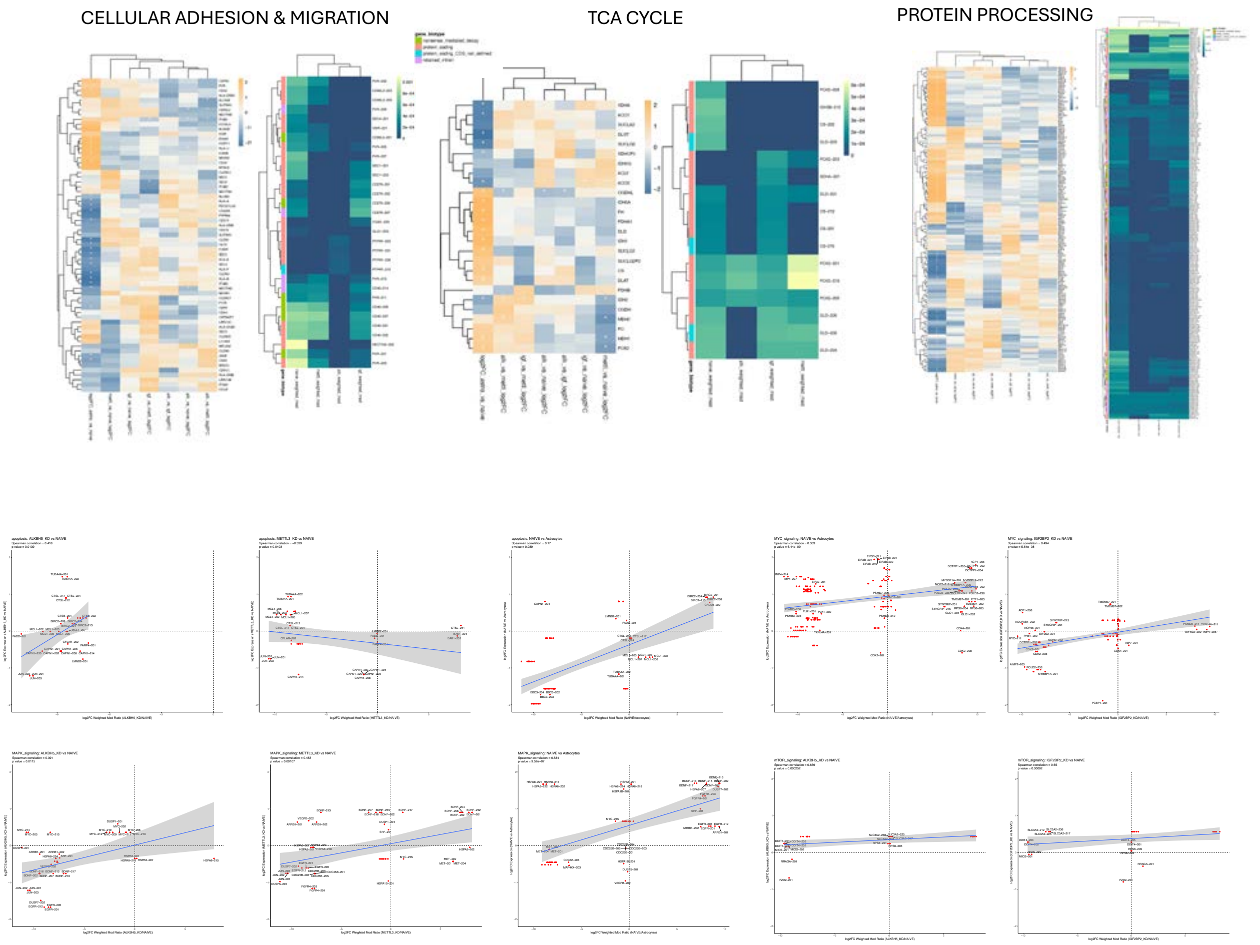


**Supplementary Figure 8: Summary of enriched pathways and the resulting effect on gene expression and methylation.** Heatmap for selected pathways where each row in a heatmap specifies a differentially expressed gene that is known to be associated with the pathway. The heatmaps depict how the methylation and gene expression can differ depending on the pathway. Correlation analysis between m6A methylation in transcripts and gene expression in Knockdown (KD) vs Naïve glioma cells for specific pathways.

#### Supplementary Figure 9

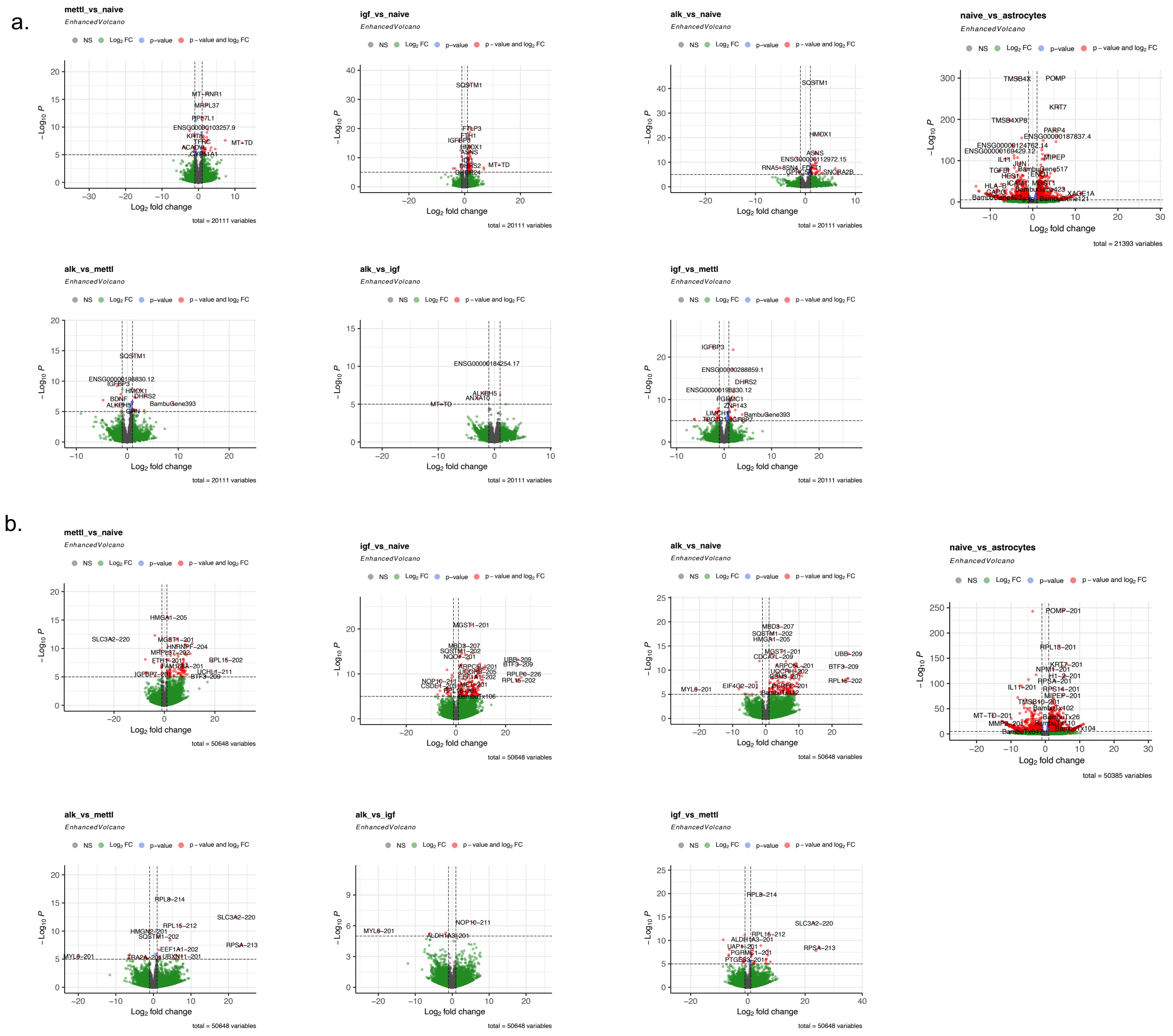


**Supplementary Figure 9: Enhanced volcano plots for the differential gene and transcript expression for Knockdown and Naïve conditions.** (a) Differential gene expression with log2 fold change greater than 0.58 or less than -0.58 highlighted in green, and significantly differentially expressed genes highlighted in red. Subset of significant genes are labelled with respective gene id. (b) Differential transcript expression with log2 fold change greater than 0.58 or less than -0.58 highlighted in green, and significantly differentially expressed genes highlighted in red. Subset of significant transcripts are labelled with respective transcript id.
